# Modeling longitudinal imaging biomarkers with parametric Bayesian multi-task learning

**DOI:** 10.1101/593459

**Authors:** Leon M. Aksman, Marzia A. Scelsi, Andre F. Marquand, Daniel C. Alexander, Sebastien Ourselin, Andre Altmann, for ADNI

## Abstract

Longitudinal imaging biomarkers are invaluable for understanding the course of neurodegeneration, promising the ability to track disease progression and to detect disease earlier than cross-sectional biomarkers. To properly realize their potential, biomarker trajectory models must be robust to both under-sampling and measurement errors and should be able to integrate multi-modal information to improve trajectory inference and prediction. Here we present a parametric Bayesian multi-task learning based approach to modeling univariate trajectories across subjects that addresses these criteria.

Our approach learns multiple subjects’ trajectories within a single model that allows for different types of information sharing, i.e. *coupling*, across subjects. It optimizes a combination of uncoupled, fully coupled and kernel coupled models. Kernel-based coupling allows linking subjects’ trajectories based on one or more biomarker measures. We demonstrate this using Alzheimer’s Disease Neuroimaging Initiative (ADNI) data, where we model longitudinal trajectories of MRI-derived cortical volumes in neurodegeneration, with coupling based on APOE genotype, cerebrospinal fluid (CSF) and amyloid PET-based biomarkers. In addition to detecting established disease effects, we detect disease related changes within the insula that have not received much attention within the literature.

Due to its sensitivity in detecting disease effects, its competitive predictive performance and its ability to learn the optimal parameter covariance from data rather than choosing a specific set of random and fixed effects a priori, we propose that our model can be used in place of or in addition to linear mixed effects models when modeling biomarker trajectories. A software implementation of the method is publicly available.

## Introduction

Despite their value in characterizing the course of neurodegeneration (Freeborough and Fox, 1997; Smith et al., 2001), repeated measures over time (i.e. longitudinal data) in neuroimaging are often limited to a baseline measurement and a small number of follow-up time-points per subject. This is primarily due to the costs and complexities of collecting such data. Consequently, within-subject trajectory models of regions of interest (ROIs) or clinical measures that are based on such limited data may not be robust to measurement errors from image acquisition or post-processing. Such noise may lead to poor inferences of true underlying trajectory parameters and poor predictions of future values, diminishing the value of trajectory based biomarkers (Curran et al., 2010). An additional problem is a limit to the flexibility of the models that can be estimated: with two time-points one can only estimate a linear model, with three only a quadratic, and so on.

There has been growing interest in methods that efficiently use longitudinal neuroimaging data; e.g. Telzer et al., (2018) provide an overview related to fMRI analysis. By far the most popular approaches are based on mixed effects modeling, which combines fixed effects, i.e. pooling subjects’ data to create an average trajectory for all subjects, with random effects, i.e. individualizing models about the average trajectory. The mixed effects modeling approach is well suited to both balanced (fixed number of samples, fixed time interval between samples) and unbalanced (varying samples or time intervals) longitudinal designs, allowing for separate analysis of between and within subject variability (Laird and Ware, 1982; Fitzmaurice et al., 2011). Bernal-Rusiel et al., (2013) and Guillaume et al., (2014) provide overviews of linear mixed effects (LME) models within neuroimaging and apply them to Alzheimer’s disease (AD).

Features from longitudinal measurements remove inter-individual differences and thus make for better descriptions of disease progression. Recent models of disease progression have integrated both cross-sectional and longitudinal information to estimate discrete or continuous disease stages for individuals (Fonteijn et al., 2012; Jedynak et al., 2012; Donohue et al., 2014; Young et al., 2014; Lorenzi et al., 2017; Schiratti et al., 2017) They have been inspired by and seek to quantify the hypothetical models of disease progression proposed by neurodegenerative disease researchers (Buckner et al., 2005; Jack et al., 2010). Oxtoby and Alexander, (2017) provide an overview of the methods within this emerging field. While the purpose of disease progression modeling is to estimate disease stage and find group-level (typically monotonic) trajectories for each biomarker, this procedure can be thought of as a form of coupling of biomarker trajectories across subjects. However, most of these models are not explicitly setup to couple subjects’ trajectories within each biomarker (e.g. a brain structural ROI) based on other biomarkers’ information (e.g. genetic risk or brain amyloid deposition). As such these models may not fully capitalize on valuable multi-modal information that may improve trajectory estimates within each biomarker.

Multi-kernel learning (MKL) presents an approach to combining multiple biomarker similarity measures. It has been previously applied to neuroimage-based pattern recognition and machine learning to discriminate disease (Hinrichs et al., 2011; Zhang et al., 2011). Young et al., (2013) applied a Bayesian MKL approach to AD discrimination which avoided the need for costly cross-validation when tuning the kernel weightings. In addition to a more efficient means of tuning such hyperparameters, Bayesian modeling also provides a principled approach to incorporating prior information, comparing models, making probabilistic predictions and inferring distributions over parameters (Gelman et al., 2013). It has been applied in many contexts within neuroimaging (Woolrich, 2012) and more specifically in both parametric and nonparametric trajectory models (Lorenzi et al., 2015; Ziegler et al., 2015).

In this paper, we develop an approach that realizes the benefits of MKL within a Bayesian trajectory model rather than a disease classification model. To do so we use multi-task learning, which aims to learn multiple related tasks simultaneously, sharing information across tasks. Bayesian MTL was previously applied in neuroimaging within the context of multi-subject fMRI analysis (Andre F. Marquand et al., 2014) based on the method proposed by Bonilla et al., (2008). Bayesian MTL has also been applied to imaging genetics via a hierarchical Bayesian model that encourages both individual and group sparsity (Nathoo et al., 2016; Greenlaw et al., 2017). Here we set the learning of each subject’s biomarker trajectory as a task and apply MTL to share information across subjects. We develop a parametric extension of Marquand et al.’s approach, a joint Bayesian linear regression that allows for full coupling across all subjects along with coupling based on biomarker similarity, so that subjects with similar measures in one biomarker may have more similar trajectories in another. Furthermore, we use MKL to couple trajectories within a biomarker based on an optimal balance of multiple other biomarkers’ information. We compare our approach, which learns the optimal parameter covariance from data to standard LME modelling, where the parameter covariance structure depends on the *a priori* choice of random and fixed effects.

This paper (i) contributes a parametric model that learns a separate trajectory for each subject while allowing for information-sharing across subjects and the integration of multi-modal information during model training, resulting in better predictions and inferences; (ii) performs simulations to validate the model and understand its properties; (iii) applies the model to clinical neuroimaging data, modeling cortical region of interest (ROI) trajectories in neurodegeneration using various biomarkers for coupling and (iv) interprets and discusses the results.

## Methodology

### Model: parametric Bayesian MTL

We present a univariate model of the temporal trajectory of a scalar variable (e.g. values of an ROI or a clinical measure) across multiple subjects. We set the learning of each subject’s trajectory as a task and use MTL to share information across subjects to better learn all subjects’ trajectories as a single, coupled model. Empirical Bayes allows us to automatically tune the degree and type of coupling across subjects using hyperparameters that control the overall covariance structure of the parameters being learned.

We start by specifying a single, large model for all *n* subjects’ trajectories, stacking the longitudinal observations of all subjects into the vector 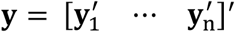, where **y**_i_ is an m_i_ × 1 vector of observations for the i^th^ subject. In total, there are 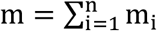 observations across subjects, so that **y** is anm × 1 vector. To model these trajectories, we can fit polynomial functions of time (e.g. age) using the following model structure:

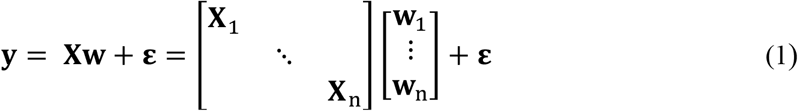

where the overall design matrix **X** is block diagonal for a chosen polynomial model of order p, with zeros in the off-diagonal entries. Defining d = p + 1, we have block **X**_i_ as the m_i_ × d design matrix for subject i having m_i_ observations at times 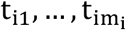:

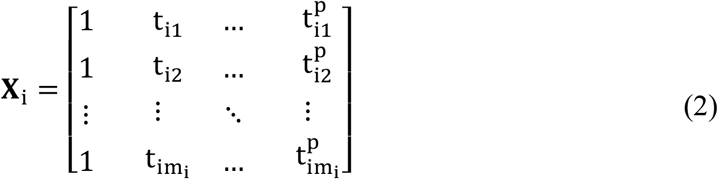

and **w** is an nd × 1 vector of parameters across subjects. If we assume additive Gaussian noise **ε** ~ N(**0**, β^−1^**I**_m_) we can solve the general linear model (GLM) formed by Eq. (1) via ordinary least squares (OLS) regression, finding a set of parameters **w**_1_, …, **w**_n_ that describe the subjects’ temporal trajectories. Each **w**_i_ is a d × 1 vector containing the trajectory parameters for subject i. In the case of linear models **w**_i_ is a 2 × 1 and contains an intercept and a slope term.

OLS regression is a simple and widely used means of modeling trajectories models for each subject. However, it assumes an independent model for each subject, thereby ignoring the similarity in other subjects’ trajectories that may greatly improve both prediction and parameter inference. Using a Bayesian approach, we propose to overcome this problem by placing a prior probability distribution over these parameters. The form of the prior we propose is:

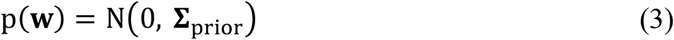

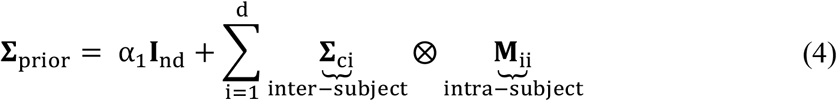

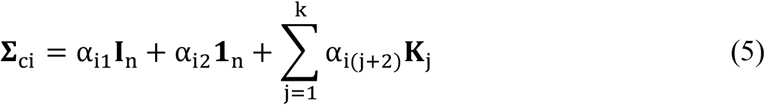

where **M**_ii_ is defined below and **Σ**_prior_ is of size nd × nd. The first term in Eq. (4), with weight α_1_, allows for a diagonal (i.e. independent) covariance structure in the parameters^1^ and ensures the matrix is positive definite. The second term is a sum of d Kronecker products. When fitting linear models (i.e. polynomial models of degree one, as we will do throughout this paper), there are two such products: one for the 0^th^ order parameters (i.e. intercepts), the other for the 1^st^ order parameters (i.e. slopes or rates of change). In each case, we take the Kronecker product of an inter-subject (coupling) matrix and an intra-subject matrix to form part of the overall covariance matrix. Each **Σ**_ci_ is an n × n matrix parametrized to allow for fully independent parameters (α_i1_ **I**_n_ term, where **I**_n_ is an n-dimensional identity matrix), fully coupled parameters (α_i2_ **1**_n_ term, where **1**_n_ is an n-dimensional matrix of ones) and coupling based on the set of k biomarker based kernels (the **K**_j_’s, each an n-dimensional positive definite matrix). The form of these biomarker kernels is detailed later in this paper. As a result, each **Σ**_ci_ contains at least k + 2 hyperparameters^2^ and overall there are at least d(k + 2) covariance-related hyperparameters. It is important to bear in mind the distinction between the hyperparameters (the α’s) used to control coupling among subjects and the parameters of individuals’ trajectory models (the **w**_i_’s stacked within **w**).

The intra-subject matrix **M**_ii_ describes how the trajectory parameters within each subject’s model are related to each other. We have chosen each **M**_ii_ to be a indicator matrix equal to one on the i-th diagonal element and zero elsewhere, so that there is no information sharing between different parameter types (e.g. between intercepts and slopes) within and across subjects. This prior structure allows us to learn the inter-subject coupling separately for each parameter type, giving the model a great deal of flexibility.

Note that simpler parametrizations of the covariance matrix are possible. For instance, choosing **Σ**_prior_ = **Σ**_c_⊗**I**_d_ with **I**_d_ the d × d identity matrix and **Σ**_c_ = α_1_**I**_n_ + α_2_**1**_n_ is closest to the form used in Marquand et al. Using instead 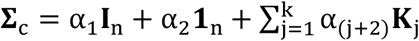 implements MKL. Both variations use fewer hyperparameters than our proposed parametrization. However, these simpler models tie the coupling of parameters types (e.g., intercepts and slopes) together and as such may be more prone to inducing spurious coupling in one set of parameters while capturing true coupling in another (e.g. spuriously coupling intercepts along with slopes). This may, in turn, lead to increased false positive group differences in subjects’ parameters.

This prior structure differs from that used in hierarchical Bayesian modeling, where individuals’ first level parameter prior means are specified as linear combinations of second level parameters that may include covariates and grouping variables, that also have prior distributions. In contrast, we assume a zero-mean prior on individuals’ parameters and incorporate covariates via kernels within each **Σ**_ci_. Kernels allow for non-linear similarity measures between covariates, adding modeling flexibility not present in linear hierarchical models. Hierarchical models, in contrast, offer a well-developed framework for modeling fixed and random effects. The two approaches are not mutually exclusive: it is possible to combine them, though for the sake of simplicity we leave this as a topic for future work.

Finally, we choose the likelihood term, i.e. the data observation model, to resemble the GLM from eq. (1), setting

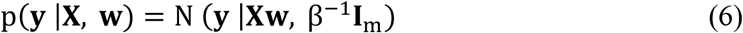

with **X** and **w** defined as before. We allow the model to learn the (inverse) measurement noise level β within the likelihood as an additional hyperparameter. With the prior and likelihood thus specified, we can use Bayes’ rule to update our beliefs on the parameter distribution given some observed data (i.e. find the posterior distribution). In this case as we have a Gaussian prior and a Gaussian likelihood the posterior is also Gaussian and has the following closed-form solution (Bishop, 2007):

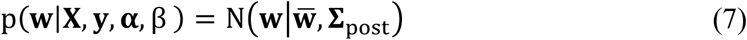

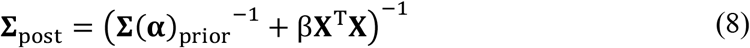

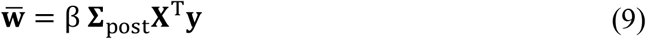

where we have collected the covariance prior’s hyperparameters into the vector **α** and have made the dependence of the prior covariance on these parameters explicit using the notation **Σ**(**α**)_prior_.

To estimate optimal values for **α** and β, we take the empirical Bayesian approach described in Huertas et al., (2017) and Marquand et al., (2014), finding the **α** and β that maximize the marginal likelihood of the observed data under our assumed model structure. With the prior as in Eq.’s (3) – (5) and the likelihood as in Eq. (6) the log marginal likelihood becomes:

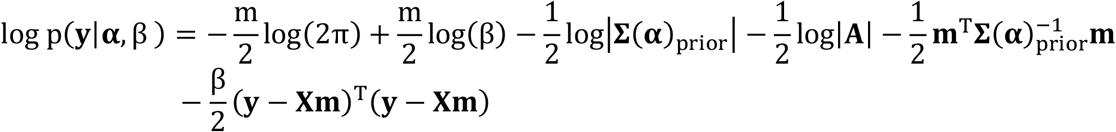

where 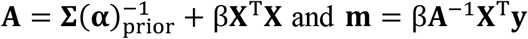 and **m** = β**A**^−1^**X**^T^**y**. We used *minimize*, a conjugate gradients optimizer available within the gpml toolbox (Rasmussen and Nickisch, 2010), which uses partial derivatives of the marginal likelihood with respect to each of the hyperparameters. We optimized these variables in the log domain to ensure positivity (see Appendix for further details).

In this paper we predict the biomarker value at each subject’s mean baseline and final follow-up ages, taking the probabilistic approach of integrating over all possible posterior model parameters. With the G−au1ssian posterior as in Eq.’s (7) – (9) and the Gaussian likelihood p(**y**_*_|**X**_*_, **w**) = N (**y**_*_ |**X**_*_**w**, β^−1^**I**_m_) of observing predictions **y**_*_ given input **X**_*_, the predictive distribution that results from integration has a closed form solution (Rasmussen, 2006):

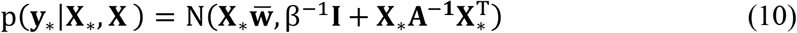

where 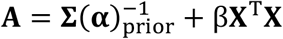 and **X**_*_ is an n × nd design matrix with a row encoding either the mean baseline age or final sample age in this case (or 2n × nd to predict both at once). In general, we can predict an arbitrary number of time-points per subject by modifying **X**_*_ accordingly.

For all models, we standardize (i.e. z-score) the training data across all subjects (the longitudinal observations **y** and each non-constant column of the design matrix **X**) as well as the out-of-sample testing data (**X**_*_, using the means and standard deviations from **X**) during model building and prediction. This ensures that all trajectories are modelled on a similar scale, which aids numerical stability when optimizing the hyperparameters. We rescale both the predictions and the estimated parameters back to their original dimensions for subsequent analysis (e.g. estimating annualized rates of change and group differences in parameters). With higher order models, the columns of **X** may be highly correlated, leading to unstable variance estimates. In such cases, one may orthogonalize the columns prior to fitting the model.

We use the log Bayes factor, a ratio of the logarithm of model evidences (i.e. marginal likelihoods) to compare biomarker-information based coupling to random-information based coupling. Log Bayes factors are a principled way of comparing Bayesian models, under the assumption that each model has the same prior probability (Penny, 2012; Penny et al., 2004).

### Software: model and figures

A MATLAB implementation of our method is available at https://github.com/LeonAksman/bayes-mtl-traj. The brain images in Figures 4, 5, 6, S7, S8 and S10 were produced via a command-line image render and snapshot tool, available at https://github.com/LeonAksman/vtkSnap.

### Simulations: data generation

We created a simulation of subjects’ trajectories that allowed us to compare several different versions of our proposed model along with a baseline model. We simulated two scenarios: (i) linear trajectories with intercept variation: group differences in intercept with fixed slope and (ii) linear trajectories with slope variation: group differences in slope with fixed intercept. In both cases, we simulated 200 subjects’ trajectories at each simulation run. To simulate intercept variation, for a given subject we randomly selected an intercept w_i0_ from the set {−10,−8,−6,−4,−2} and a fixed slope of w_i1_ = −1, and then randomly selected an initial measurement time t_i1_ between 0 and 10. We then generated three simulated samples with fixed time intervals: y(t_i1_) = w_i0_ + w_i1_t_i1_ + *ε*_i1_ = w_i0_ − t_i1_ + *ε*_i1_, y(t_i2_ = t_i1_ + 0.05) = w_i0_ − t_i1_ − 0.05 + *ε*_i2_ and y(t_i3_ = t_i1_ + 0.10) = w_i0_ − t_i1_ − 0.10 + *ε*_i3_, where *ε*_i1_, *ε*_i2_, *ε*_i3_ are three independent measurement errors, each drawn from N(0, σ_m_). We simulated four different levels of measurement noise, σ_m_ = 1,2,4,8. For each noise level, we made thirty simulation runs and evaluated nine different models (described below) on each run. We used the first two samples of each subject (t_i1_, t_i2_ for subject i) to train each model and the third sample (t_i3_ for subject i) to evaluate out-of-sample prediction accuracy. The top row of Figure 1 provides an example of one simulation run for 200 subjects at each noise level.

To simulate slope variation, for a given subject i we randomly selected a slope w_i1_ from the set {−1.0,−1.5,−2.0,−2.5,−3.0} and a fixed intercept of w_i1_ = 0. We randomly selected t_i1_ as before and generated y(t_i1_), y(t_i2_ = t_i1_ + 0.05) and y(t_i3_ = t_i1_ + 0.10) with measurement errors *ε*_i1_, *ε*_i2_, *ε*_i3_ drawn from N(0, σ_m_). We used the same four measurement noise levels and again made thirty simulation runs, generating 200 simulated subjects’ trajectories for each noise level. We evaluated the same nine models using the first two samples of each subject for training and the third for prediction. The bottom row of Figure 1 provides an example of one simulation run at each noise level.

**Figure 1.**
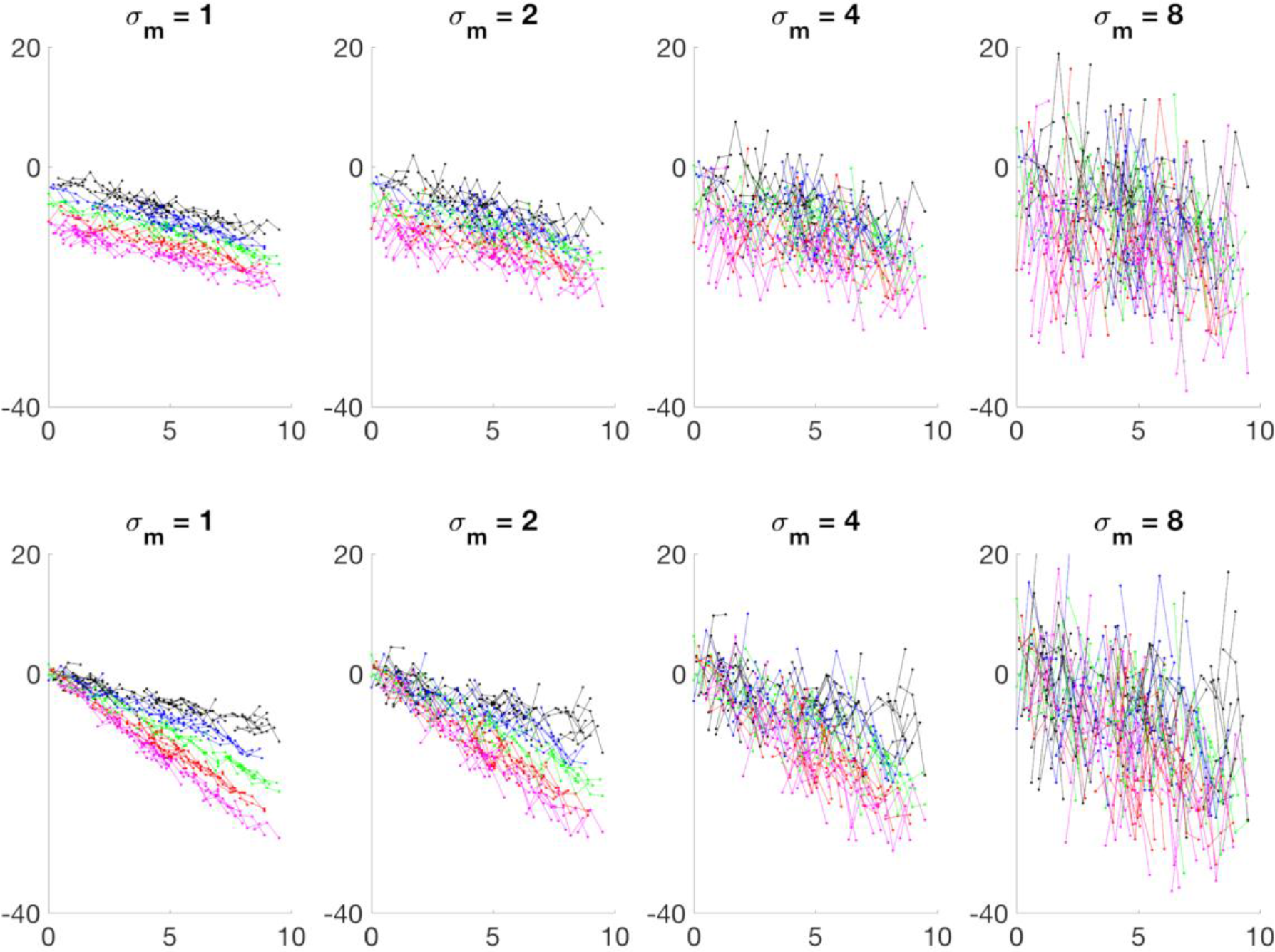
**Top row:** one run of a simulation of 200 subjects’ longitudinal samples with group differences in intercept at four different measurement noise (*σ*_*m*_ levels. **Bottom row:** the same with group differences in slope. Each subject has three samples, with trajectory parameters chosen from among five gradations of intercept (top row) or slope (bottom row), indicated by different colors.

### Simulations: model building

We investigated how different variations of the model, with and without biomarker-kernel-based coupling (i.e. via the **K**_j_ matrices described above), performed under the two parameter variation scenarios and the four measurement noise levels. We created a 200 × 1 biomarker vector **b** as a noisy measure of the group difference in the parameters, so that the i^th^ element of **b** was b_i_ = w_i0_ + *ε*_i_ under intercept variation and b_i_ = w_i1_ + *ε*_i_ under slope variation, with *ε*_i_ drawn from N(0, 1) in both cases^3^. Using this biomarker, we comT pared two commonly used similarity matrices: (i) a rank-one approximation **K**(**b**)_linear_ = **bb**^T^ (the outer product of **b** with itself); and (ii) a squared exponential (SE) kernel (also referred to as a Gaussian radial basis function kernel) **K**(**b**)_SE_, with k_ij_ = exp (−σ_SE_‖b_i_ − b_j_‖^2^) in the i^th^ row and j^th^ column, where we make the dependence of these matrices on the input explicit. The term σ_SE_ > 0 is a parameter that gives the kernel additional scaling flexibility. It is also possible to encode categorical (i.e. group) membership via a binary similarity matrices rather than an SE kernel, with one indicating two subjects belong to the same class and zero otherwise.

When using SE kernels, we treated the σ_SE_ term as a covariance prior hyperparameter (in addition to the α’s in Eq.’s (4) and (5)). We formed six different models using these two kernels: ‘*linear both*’ and ‘*Gaussian both*’ had covariance prior structure as in Eq.’s (4) and (5), parameterizing full independence, full coupling and kernel-based coupling in both the intercept and slope parameters. The ‘*linear int*’ and ‘*Gaussian int*’ models restricted kernel-based coupling to the intercepts: referring to Eq.’s (4) and (5) and assuming one coupling kernel **K**_1_, this means we allow a **K**_1_ term in **Σ**_c1_ but not in **Σ**_c2_. The ‘*linear slope*’ and ‘*Gaussian slope*’, in contrast, restricted kernel-based coupling to the slopes, allowing a **K**_1_ term in **Σ**_c2_ but not in **Σ**_c1_. These latter four models allowed us to test the effect of ‘oracle-like’ (i.e. with perfect *a priori*) knowledge of simulation scenario: e.g. whether the two ‘*int*’ models outperform other models in the variable intercept, fixed slope scenario. In such cases, biomarker-based coupling of slopes in the intercepts variation case (or vice versa) is extraneous and may lead to spurious inference of group differences if a model is allowed to infer coupling where none exists.

We compared biomarker coupled models to several simpler coupled models. The simplest of these was the ‘*OLS*’ model, an uncoupled model which asymptotically corresponds to a Bayesian model with a parameter covariance prior of *α*_1_**I**_2n_ with *α*_1_ tending to infinity (i.e. a high prior uncertainty on all parameters for all subjects). The second model, ‘*plain*’, trades off fully independent and fully coupled covariance priors, without any kernel-based coupling. It is very similar to an LME model with random intercepts and random slopes. To understand the role of kernel-based coupling, we compared biomarker-kernel coupled models to random-information-kernel coupled models. We formed a 200 × 1 vector **r** with each element drawn from N(0, 1), so that each subject was assigned a random number, and formed another SE kernel **K**(**r**)_SE_ based on it. We used this to create ‘*random*’, parametrized exactly as ‘*Gaussian both*’.

We compared our approach to standard LME modelling using the LME implementation available in Freesurfer (Bernal-Rusiel et al., 2013; Bernal-Rusiel et al., 2013)^4^. Specifically, we built two LME models, with fixed effects of baseline age and baseline biomarker value and either: random intercepts (termed ‘*LME: rI*’) or random intercepts and slopes (‘*LME: rI, rS*’).

We also compared our empirical Bayesian approach, which produces point estimates of hyperparameter priors (i.e. an estimate of prior means with zero variance) to a fully Bayesian approach, in which we place priors on the hyperparameters and estimate their posterior distribution. In this way, the fully Bayesian approach accounts for the hyperparameter uncertainty, which may improve parameter and prediction coverage by improving the estimation of their uncertainties. We compared the empirical Bayesian version of ‘*plain*’, with five covariance hyperparameters (the **α**′s) and one inverse observation noise parameter (β to a fully Bayesian model with the same covariance structure. We placed broad, uninformative half-normal priors on the **α**’s, by setting each α ~ normal(0,100) with constraint α > 0, and an inverse Gamma distribution on β^−1^, setting β^−1^~InvGamma(1,1). We used Markov chain Monte Carlo (MCMC) to estimate the full model as it was no longer analytically tractable to derive all the necessary posterior distributions. We implemented the full model in Stan (Carpenter et al., 2017) using Hamiltonian Monte Carlo. Due to the significantly longer running times of the full model (see Results) we ran the same two simulation scenarios for 50 instead of 200 subjects, with all other settings as before. We used the default parameters for MCMC sampling: four chains, with 1000 warmup iterations and 1000 sampling iterations per chain, so that the posterior distributions had 4000 sampling iterations in total. We checked the convergence of the chains’ posterior distributions using the metric provided. We checked the convergence of the chains’ posterior distributions using the 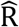 metric provided. We checked the quality of the chain by looking at the means, Markov Chain standard errors (MCSE) and effective sample sizes of the hyperparameters.

Finally, we performed additional simulations to test our assumption of independent measurement noise, parameterized by β in Eq. (6), by instead allowing a block diagonal noise structure. For each subject’s measurements, we used a noise covariance with 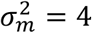 on the diagonal and all off-diagonal terms set to 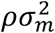, so that we recover uncorrelated noise when *ρ* = 0 and perfectly correlated noise (i.e. the same for all observations over time) when *ρ* = 1. We simulated with three levels of *ρ* (0, 0.5, 0.75).

**Table 1.** Models fit at each simulation run. Last column contains number of hyperparameters in given covariance prior.

| Model | Purpose | Kernel | Covariance Prior | # Cov. Hyper’s |
| --- | --- | --- | --- | --- |
| *random* | Allow both slope and intercept coupling via random biomarker | $\mathbf{K}(\mathbf{r})_{\text{SE}}$ | $\alpha_1 \mathbf{I}_{2n} + \sum_{i=1}^2 (\alpha_{i1} \mathbf{I}_n + \alpha_{i2} \mathbf{1}_n + \alpha_{i3} \mathbf{K}(\mathbf{r})_{\text{SE}}) \otimes \mathbf{M}_{ii}$ | 9 |
| *linear both* | Allow both slope and intercept coupling via true biomarker | $\mathbf{K}(\mathbf{b})_{\text{linear}}$ | $\alpha_1 \mathbf{I}_{2n} + \sum_{i=1}^2 (\alpha_{i1} \mathbf{I}_n + \alpha_{i2} \mathbf{1}_n + \alpha_{i3} \mathbf{K}(\mathbf{b})_{\text{linear}}) \otimes \mathbf{M}_{ii}$ | 7 |
| *Gaussian both* | Allow both slope and intercept coupling via true biomarker | $\mathbf{K}(\mathbf{b})_{\text{SE}}$ | $\alpha_1 \mathbf{I}_{2n} + \sum_{i=1}^2 (\alpha_{i1} \mathbf{I}_n + \alpha_{i2} \mathbf{1}_n + \alpha_{i3} \mathbf{K}(\mathbf{b})_{\text{SE}}) \otimes \mathbf{M}_{ii}$ | 9 |
| *linear int* | Allow intercept coupling via true biomarker | $\mathbf{K}(\mathbf{b})_{\text{linear}}$ | $\alpha_1 \mathbf{I}_{2n} + (\alpha_{11} \mathbf{I}_n + \alpha_{12} \mathbf{1}_n + \alpha_{13} \mathbf{K}(\mathbf{b})_{\text{linear}}) \otimes \mathbf{M}_{11} + (\alpha_{21} \mathbf{I}_n + \alpha_{22} \mathbf{1}_n) \otimes \mathbf{M}_{22}$ | 6 |
| *Gaussian int* | Allow intercept coupling via true biomarker | $\mathbf{K}(\mathbf{b})_{\text{SE}}$ | $\alpha_1 \mathbf{I}_{2n} + (\alpha_{11} \mathbf{I}_n + \alpha_{12} \mathbf{1}_n + \alpha_{13} \mathbf{K}(\mathbf{b})_{\text{SE}}) \otimes \mathbf{M}_{11} + (\alpha_{21} \mathbf{I}_n + \alpha_{22} \mathbf{1}_n) \otimes \mathbf{M}_{22}$ | 7 |
| *linear slope* | Allow slope coupling via true biomarker | $\mathbf{K}(\mathbf{b})_{\text{linear}}$ | $\alpha_1 \mathbf{I}_{2n} + (\alpha_{11} \mathbf{I}_n + \alpha_{12} \mathbf{1}_n) \otimes \mathbf{M}_{11} + (\alpha_{21} \mathbf{I}_n + \alpha_{22} \mathbf{1}_n + \alpha_{23} \mathbf{K}(\mathbf{b})_{\text{linear}}) \otimes \mathbf{M}_{22}$ | 6 |
| *Gaussian slope* | Allow slope coupling via true biomarker | $\mathbf{K}(\mathbf{b})_{\text{SE}}$ | $\alpha_1 \mathbf{I}_{2n} + (\alpha_{11} \mathbf{I}_n + \alpha_{12} \mathbf{1}_n) \otimes \mathbf{M}_{11} + (\alpha_{21} \mathbf{I}_n + \alpha_{22} \mathbf{1}_n + \alpha_{23} \mathbf{K}(\mathbf{b})_{\text{SE}}) \otimes \mathbf{M}_{22}$ | 7 |
| *plain* | No biomarker based coupling | none | $\alpha_1 \mathbf{I}_{2n} + (\alpha_{11} \mathbf{I}_n + \alpha_{12} \mathbf{1}_n) \otimes \mathbf{M}_{11} + (\alpha_{21} \mathbf{I}_n + \alpha_{22} \mathbf{1}_n) \otimes \mathbf{M}_{22}$ | 5 |
| *OLS* | No coupling | none | $\alpha_1 \mathbf{I}_{2n}, \alpha_1 \rightarrow \infty$ | 0 |

### Simulations: model evaluation

Parametric Bayesian modeling provides probabilistic predictions (via Eq. (10)) and parameter distributions (via Eq.’s (7) – (9)). We can thus compare the performance of models in terms of both their accuracy of predicting ground truth trajectories and how well they estimate model parameters. The latter objective is potentially important when there are group differences in trajectories, e.g. when a disease group has a steeper rate of gray matter volume decline in a region of interest (ROI) compared to a healthy control group. In such a case, using linear models we should be able to detect a difference in the slope parameters across the groups and, assuming the decline starts from the same level, no corresponding difference in intercepts.

For each model, we evaluated the mean absolute error (MAE) of predicting subjects’ held-out samples and quantified the accuracy of the inferred trajectory parameters (intercepts and slopes) via coverage probability and MAE measures. We defined the coverage probability as the fraction of times the true value of a parameter (intercept or slope) falls within two standard deviations of its estimated value, i.e. within the posterior credible interval of the parameter. As this is a 95% credible interval a coverage probability of 0.95 is an ideal outcome. For the Bayesian MTL models, we can easily calculate these quantities using the posterior means and variances (Eq.’s (7) – (9)). For the LME models, direct estimates of the posterior parameter variance were not available: we therefore estimated them by adding the fixed effect and random effect variance estimates of the intercept and slope when appropriate.

In practice, the coverage probability may not be sufficient for understanding how accurately a model estimates a parameter as a model may simply estimate a high enough variance so that the true value is always covered. For this reason, we also computed the error (MAE) between the estimated parameters (intercept, slope) and their known true values.

### ADNI application: dataset

Data used in the preparation of this article were obtained from the Alzheimer’s Disease Neuroimaging Initiative (ADNI) database (adni.loni.usc.edu). The ADNI was launched in 2003 as a public-private partnership, led by Principal Investigator Michael W. Weiner, MD. The primary goal of ADNI has been to test whether serial magnetic resonance imaging (MRI), positron emission tomography (PET), other biological markers, and clinical and neuropsychological assessment can be combined to measure the progression of mild cognitive impairment (MCI) and early Alzheimer’s disease (AD). For up-to-date information, see www.adni-info.org.

We used ADNI subjects with at least one [^18^F]-Florbetapir PET scan, which images brain amyloid accumulation, using the earliest available image as the baseline time-point. We chose a subset of 437 subjects: 104 cognitively normal (CN), 243 mild cognitive impairment (MCI), 90 probable Alzheimer’s disease (AD). Each had at least three MRIs at or after the chosen baseline, with a total of 1545 images across all subjects. There were 257 subjects with three MRIs, 138 subjects with four MRIs, 31 subjects with five MRIs, ten subjects with six MRIsand one subject with seven MRIs.

We processed the PET scans to derive standardized uptake value ratio (SUVR) values in cortical grey matter as measures of cortical amyloid deposition at baseline, defined here as the first Florbetapir scan available. Details of PET image processing can be found in Scelsi et al., (2018). Briefly, tracer uptakes in the cortical ROI were standardised to the uptake in a composite reference region following recommendations from (Landau et al., 2015). We also used amyloid-β, total tau and phosphorylated tau (pTau) from CSF measured at or before baseline as measures of severity of amyloid and tau pathology. Lastly, we retrieved subjects’ Apolipoprotein E (APOE) genetic information, particularly the number of ε2 and ε4 alleles (Harold et al., 2009; Liu et al., 2013).

We parcellated the T1-weighted images using geodesic information flows (GIF; Cardoso et al., 2015), creating twenty cortical sub-lobe volumes from each image (see Figure S6 in Supplementary Material for the list of ROIs). We then normalized each of these ROIs using each subject’s total intracranial volume (TIV). Normalized ROIs were subsequently used for longitudinal trajectory modeling and out-of-sample prediction. We withheld the final follow-up ROI from each model to test the out-of-sample prediction accuracy of our models.

### ADNI application: model building

We built eight different types of models, detailed in Table 2, for each of the twenty regions for a total of 160 models.

We used first order (linear) polynomial models for all regions: previous work has shown this is a reasonable assumption for modeling cortical trajectories (Ziegler et al., 2015). Based on our simulations (see Results), we chose the ‘*Gaussian both*’ type of model when using biomarker coupling, allowing both intercept and slope coupling, assuming no prior knowledge of the type of coupling that exists in the data. We formed four different kernels (**K**_1_, **K**_2_, **K**_3_, **K**_4_) based on true biomarkers along with a fifth kernel based on a random biomarker-based kernel (**K**_5_ = **K**(**r**)_SE_, **r** a vector of random values) as in the simulations. Kernels **K**_i_ = **K**(**b**_i_)_SE_, i = 1,2,3 were formed using: (i) **b**_1_, a vector of SUVR values across subjects derived from amyloid PET, (ii) **b**_2_ = log(**tau/Aβ**), a vector encoding the relationship between CSF tau, which increases in subjects with AD, and CSF amyloid-β, which decreases in those with AD (Sunderland et al., 2003), log transformed to improve normality and (iii) **b**_3_, a vector encoding CSF pTau (Hampel et al., 2010).

To encode the APOE genetic similarity between subjects we used the weighted identity by state (weighted-IBS) kernel function as in Kwee et al., (2008):

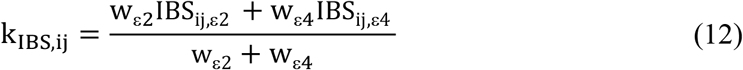

where the IBS_ij,ε2_,IBS_ij,ε4_ terms (each taking values 0, 1 or 2) refer to the number of ε2 and ε4 alleles shared by subjects i and j. The inverse minor allele frequency (1/MAF) weights w_ε2_,w_ε4_ serve to up-weight rarer SNPs. The range of this function is between zero and two. To better compare to the other SE kernels, we created an exponentiated version of this kernel function:

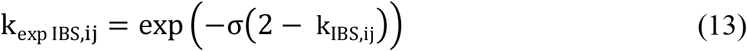

that includes a scaling hyperparameter σ and has a range between zero and one. We formed the **K**_4_ kernel matrix using this kernel function.

We also compared our approach to (Freesurfer-based) LME models. We built three LME models with fixed effects of age and baseline amyloid load (measured via PET SUVR, as in the ‘*PET amyloid*’ MTL model) and either: random intercepts (termed ‘*Rand Int*’), random intercepts and slopes (‘*Rand Int/Slp*’) or random intercepts, random slopes and random amyloid (‘*Rand Int/Slp/Amyloid*’).

**Table 2.**
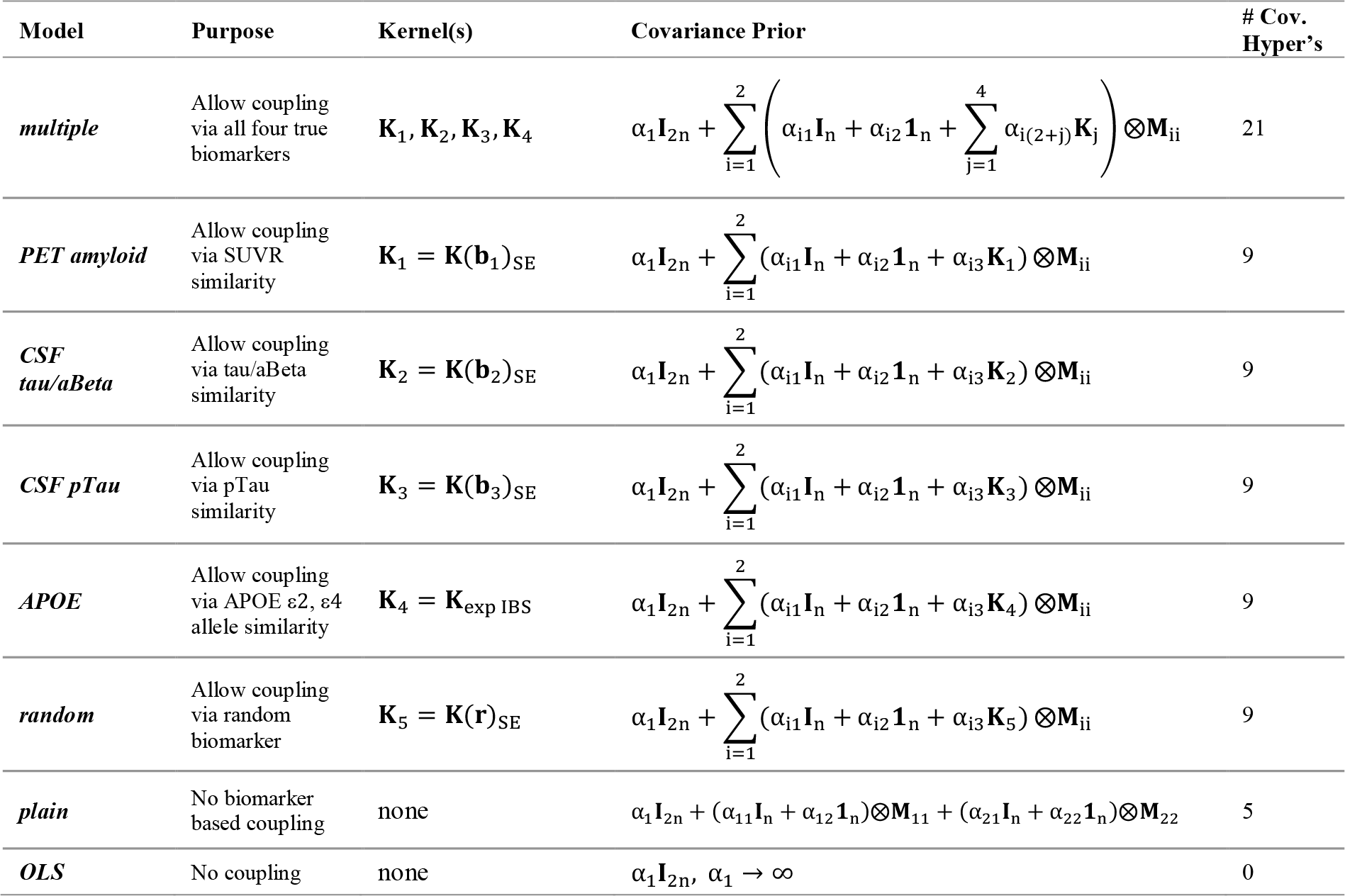
Models fit for ADNI data. Last column contains number of hyperparameters in given covariance prior.

## Results

### Simulations: results

Figure 2 depicts boxplots of the prediction MAEs across simulation runs. Models using SE kernel-based coupling (‘*Gaussian*’ type models) generally performed better than their linear kernel counterparts (‘*linear*’ type models). The advantage of the SE kernel in some cases may be attributable to the ability to tune the kernel width (the σ_SE_ term) as an additional hyperparameter, which adds scale flexibility. ‘*Gaussian both*’ was consistently among those with lowest MAE. We expected the oracle-like models (‘*int*’ type in top row, ‘*slope*’ type in bottom, marked with asterisks in the figures) to outperform the other models, however, overall, they perform similarly to the other models in most cases. Importantly, all MTL based models (including ‘*plain’*) outperform ‘*OLS*’ by a large margin, roughly halving the error. Figure S1 depicts histograms of parameter estimates for both ‘*plain*’ and ‘*OLS*’ for a representative simulation run, showing that the Bayesian model shrinks both the slopes and intercepts to their respective group mean, decreasing the variance of the estimates considerably. The shrinkage also results in a small increase in bias, evidenced by the larger distance between the parameter means of ‘*plain*’ (dashed red line) to the true parameter means (dashed black line) compared to ‘*OLS*’ (dashed blue line), with an overall large decrease in the mean squared errors of the parameter estimates (‘*plain*’: 8.8 for intercepts, 0.4 for slopes; ‘*OLS*’ 193.0 for intercepts, 8.1 for slopes).

**Figure 2.**
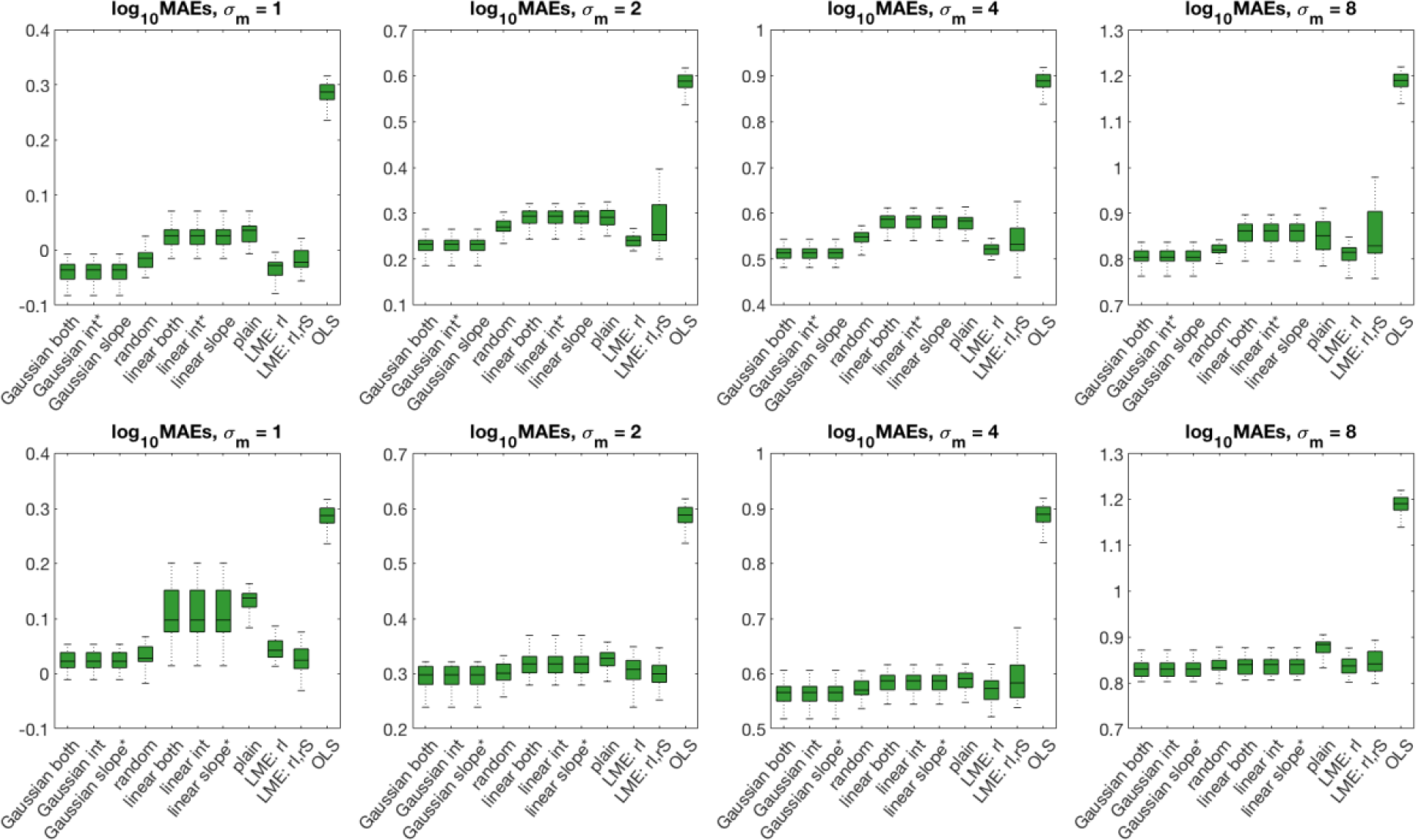
Boxplots of log mean absolute errors (MAEs) of predictions of all models across all simulations runs for the two scenarios: intercept variation (top row) and slope variation (bottom row) for four levels of measurement noise (*σ*_*m*_). Models with oracle-like prior information are marked with an asterisk (top row: ‘*int*’ models; bottom row: ‘*slope*’ models).

The two LME models also performed well, with similar MAEs to the ‘*Gaussian*’ models. Supplementary Figure S2 depicts the corresponding prediction coverage probabilities, showing that both the MTL and LME models’ predictions cover the true target value at close to the ideal rate of 0.95, again outperforming ‘*OLS*’ by a large margin, especially at higher noise levels. We also observe that the simpler MTL models (‘*linear*’ models, along with ‘*plain*’) have both high coverage in Figure S2 and relatively high MAE in Figure 2, meaning that, compared to the other MTL and LME models, they make relatively inaccurate predictions but estimate high enough measurement uncertainty to cover the true target value.

Figure 3 depicts the corresponding parameter coverage probabilities and estimation errors. In the fixed slope, varying intercepts scenario (top row), coverage of the true fixed slope parameter was high for all models (at or near 100%, exceeding the nominal level of 95%) while intercept coverage varied greatly across models and noise levels. The LME models generally did not cover the true intercept values as frequently as the MTL models, particularly the random intercepts model (‘*LME: rI*’). The random intercepts, random slopes model (‘*LME: rI, rS*’) performed better at higher noise levels, but was generally outperformed by the MTL models. One possible explanation is that the MTL models explicitly model parameter uncertainty as part of their Bayesian formulation, while we have had to estimate the LME models’ overall parameter uncertainty by combining the associated fixed and random effect uncertainties. In addition, the LME models also have higher parameter estimation errors (the intercept and slope log MAE figures in the top row), which measures the quality of parameter mean estimates rather than variances. Overall in this scenario the ‘*Gaussian*’ models outperformed all others in terms of parameter coverage and estimation error while the ‘*linear*’ models and ‘*plain*’ were comparable to the LME models in terms of parameter estimation error.

**Figure 3.**
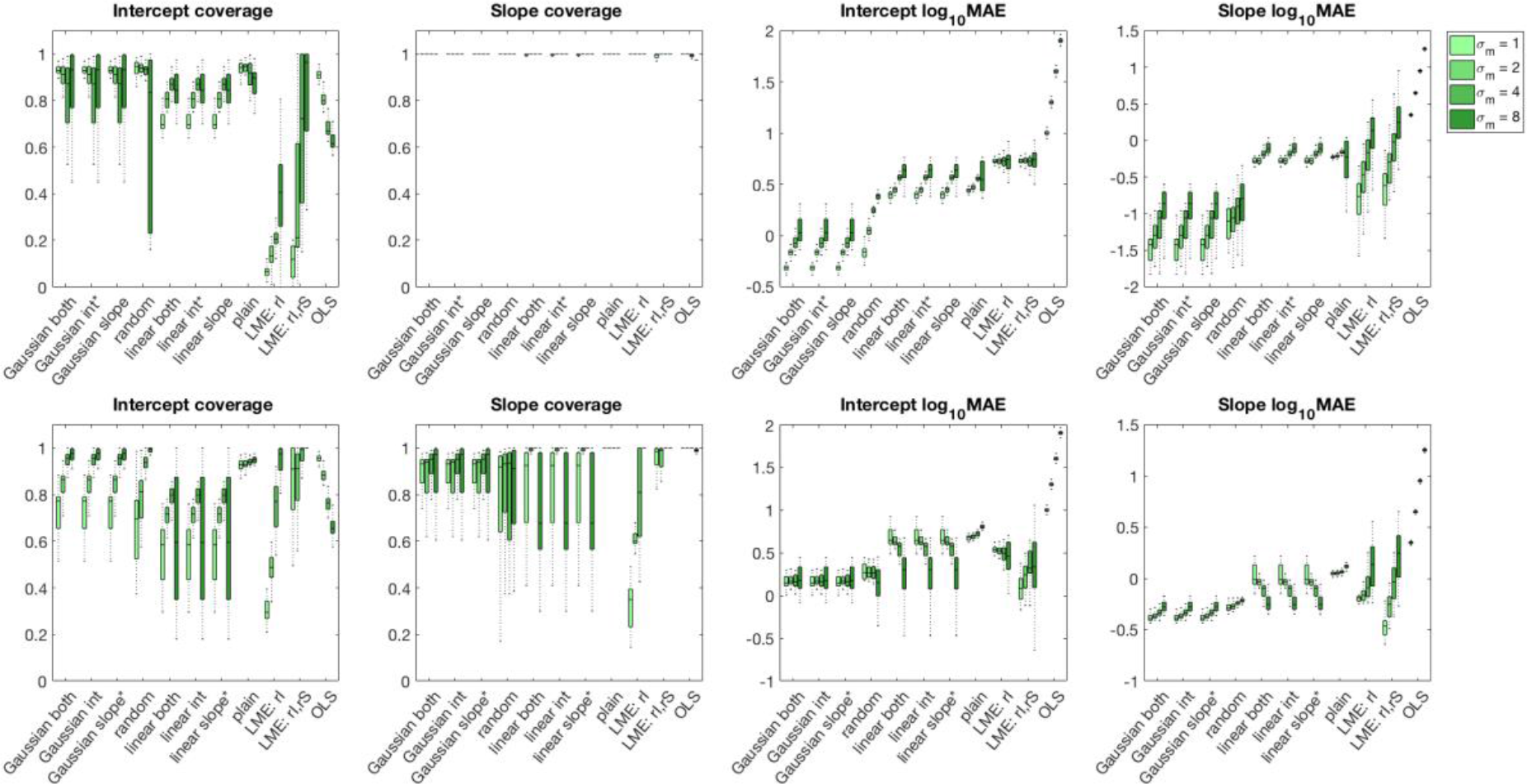
Boxplots of parameter coverage probabilities (i.e. fractions of times the true parameter value fell within the posterior credible region) and log mean absolute errors (MAE) between estimated and actual parameters for the two scenarios: intercept variation (top row) and slope variation (bottom row) for four measurement noise levels (*σ*_*m*_). Models with oracle-like prior information are marked with an asterisk (top row: ‘*int*’ models; bottom row: ‘*slope*’ models).

In the second scenario, fixed intercept and varying slopes (Figure 3, bottom row), the ‘*Gaussian*’ models performed well in both parameter coverage and parameter estimation error; in this case, the two LME models also performed competitively. ‘*LME: rI, rS*’ had consistently highest intercept and slope coverage and lowest parameter estimation errors at low measurement noise levels, reflecting the fact that the random slopes assumption is appropriate in this scenario. However, this model’s parameter estimation error, particularly the slope MAE, increases sharply with higher measurement noise levels while the ‘*Gaussian*’ models are relatively unaffected.

The two simulation scenarios suggest that the ‘*Gaussian*’ style MTL models are a good choice for both prediction and parameter inference and compare favorably with standard LME models in many cases. Among these, ‘*Gaussian both*’ is appealing as it makes no *a priori* assumptions on the type of coupling that exists within the data. Therefore, we used this type of model throughout our experiments with the Alzheimer’s study data.

Figure S3, part A in the supplementary material shows the empirical Bayesian implementation of ‘*plain*’ (‘*EB plain*’) has very similar predictive performance to the full Bayesian implementation (‘*MCMC plain*’) in terms of prediction error. Both have high coverage of the true target values; the full Bayesian model is consistently near the optimal coverage of 0.95 while the empirical Bayesian model is prone to overestimating the predictive uncertainty, so that it covers the target in most cases. Figure S3, part B depicts the parameter estimation metrics: the full Bayesian model has much higher coverage of the intercept in both scenarios; both models have similarly high coverage of the slope. The full Bayesian model has lower error in estimating the true values of both intercepts and slopes in both scenarios. We briefly compared the computation times of the two models on a single run of the intercept varying scenario with 20, 50 and 100 subjects: the empirical Bayesian model took 0.13, 0.38 and 0.50 seconds respectively while the full Bayesian model was considerably slower: 51, 398 and 4356 seconds respectively. Table S1 in the supplementary material gives the convergence diagnostics for the posterior estimates of the hyperparameters of ‘*MCMC plain*’ for one run of the intercept varying scenario. The estimates appear to have converged: all 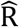 values were at their ideal values of one, the number of effective samples (N_eff_) was high in all cases and the MCSE’s were small compared to the estimated posterior means, so that the 95% confidence intervals on the means did not cross zero.

We simulated varying amounts of noise correlation across observations for each subject. We chose five representative models: ‘*Gaussian both*’ and ‘*plain*’ MTL models plus the two LME models and ‘*OLS*’. Figure S4, part A, shows that prediction metrics are similar between ‘*Gaussian both*’ and the two LME models in both scenarios. All models’ coverage increases and error falls as noise correlation increases. Figure S4, part B in the supplementary material shows corresponding parameter estimation metrics for both scenarios. ‘*Gaussian both*’ outperforms the LME models on intercept coverage and parameter error in the intercept varying scenario (top row) but ‘*LME: rI, rS*’ has near optimal coverage in the slope varying scenario (bottom row), where the random slopes assumption is appropriate. However, these two models are similar in terms of slope coverage and both intercept and slope estimation errors. Again, across the two simulation scenarios ‘*plain*’ covers the true targets by overestimating uncertainty while ‘*OLS*’ performs poorly in terms of prediction and parameter error.

### ADNI application: results

The likelihood term in Eq. (6) assumes that observations are normally distributed about their mean (i.e. the residuals are normal) and uncorrelated over time within each subject. We tested the impact of these assumptions on ADNI modelling by comparing the histograms of residuals for two models in Figure S5: ‘*CSF tau/aBeta*’, which was representative of the biomarker-coupled MTL models and ‘*OLS*’, the uncoupled reference model. Across all regions, the residuals of ‘*CSF tau/aBeta*’ are much closer to being normally distributed that those of ‘*OLS*’. We also tested for heteroscedasticity, calculating the correlation of residuals to baseline age across subjects and found no significant correlation in both models for any region.

Figure 4 (top part) depicts the true and estimated annualized rates of change across the twenty cortical ROIs for four representative models (‘*OLS*’, ‘*plain*’, ‘*random*’, ‘*multiple*’). Note that there is no information exchange between ROIs; in its presented form our method is univariate, coupling across subjects within each ROI and modeling ROIs separately. We computed the true annualized rate of change by dividing the percentage change from baseline to final (held-out) follow-up by the number of elapsed years. We note that this true annualized rate is essentially a two-point OLS estimate and is therefore more a silver than a gold standard. We see most of the cortex degenerating by 0.33% (middle cingulate) to 1.3% (posterior insula) annually, with the lateral regions generally degenerating faster than the medial regions We computed the models’ estimated annualized rates using their predictions of the held-out sample instead of the true held-out value. Figure 4 (bottom) depicts the associated MAEs of these predictions, with the three MTL models (‘*plain*’, ‘*random*’, ‘*multiple*’) having lower MAEs than the ‘*OLS*’ model across most ROIs. The two kernel coupled models (‘*random*’ and ‘*multiple*’) have further decreased MAEs compared to ‘*plain*’, though there is no further discernible difference in MAE between the two.

**Figure 4.**
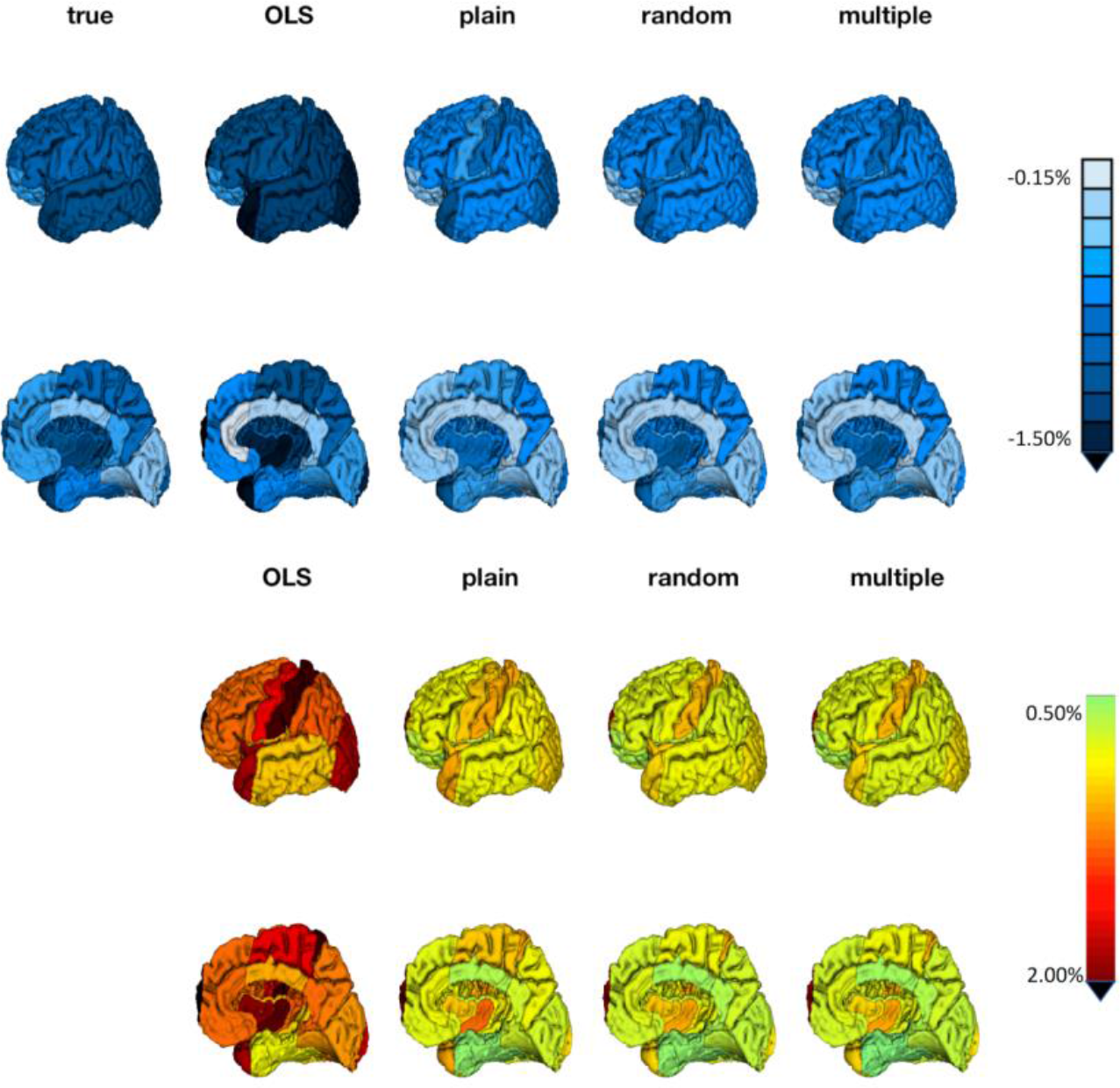
**Top:** True and estimated annualized rates of change across cortex for four representative MTL models **Bottom:** MAEs of estimates.

Figure S6 in the Supplementary Material provides a quantitative comparison of prediction error across all OLS plus MTL models. We performed t-tests on the difference in absolute error between models to understand the effect of various modeling choices, showing: (i) all MTL model errors are significantly lower than those of ‘*OLS*’ across all ROIs bar the lateral temporal region (where only ‘*plain*’ and ‘*random*’ have higher error than ‘*OLS*’); (ii) there is further improvement due to kernel coupling, evidenced by significantly lower absolute errors in at least one model relative to ‘*plain*’ in 13 out of 20 ROIs; (iii) as in simulations, there is a small difference in error between random-information-based and biomarker-based kernel coupling, with some biomarker-based models having significantly lower error than ‘*random*’ (within ten ROIs: anterior insula, DLPFC, lateral occipital, lateral parietal, lateral temporal, medial parietal, medial temporal, posterior cingulate, supratemporal and temporal pole regions). Statistical tests were Bonferroni corrected for 320 (8 models × 20 ROIs × 2 parameter types) comparisons. Furthermore, Figure S7 in the Supplementary Material depicts the MAEs of predicting annualized rate of change for three LME models (described in Methods) built using the same information as in ‘*PET amyloid*’. All models had very similar MAEs across ROIs in this case.

Figure 5 depicts log Bayes factors across cortical regions for the comparison of the five biomarker-coupled MTL models to ‘*random*’, showing ‘*CSF tau/aBeta*’, ‘*PET amyloid*’ and ‘*multiple*’ have the largest and most widespread improvements in model evidence. We also see that ‘*multiple*’ is most similar to ‘*PET amyloid*’, the best individual biomarker coupled model in terms of model evidence, providing some assurance that combining kernels works as expected.

Figure 6 depicts the significance of diagnostic group differences (CN, MCI or AD) in subjects’ estimated parameters across OLS, LME and MTL assessed via one-way ANOVAs and Bonferroni corrected for 480 (12 model comparisons × 20 ROIs × 2 parameter types). The three models with highest model evidence are depicted in Figure 6 (‘*CSF tau/aBeta*’, ‘*PET amyloid*’, ‘*multiple*’); Figure S8 depicts all MTL models, with Bonferroni correction for 320 comparisons. In both figures, cross-sectional differences in predicted volumes at mean baseline age across subjects (73.5 years) are depicted instead of intercepts. Intercepts represent group differences at age zero (i.e. at birth) while we have measured and modelled cortical degeneration in older adults. The three MTL models in Figure 6 agree that there are significant cross-sectional disease-related differences in volumes across the cortex, with sparing of the sensorimotor and cingulate regions (Figure 6, top row). The three LME models, in contrast, detect a less widespread and less significant pattern of cross-sectional differences than the MTL models while ‘*OLS*’ detects even fewer cross-sectional differences.

**Figure 5.**
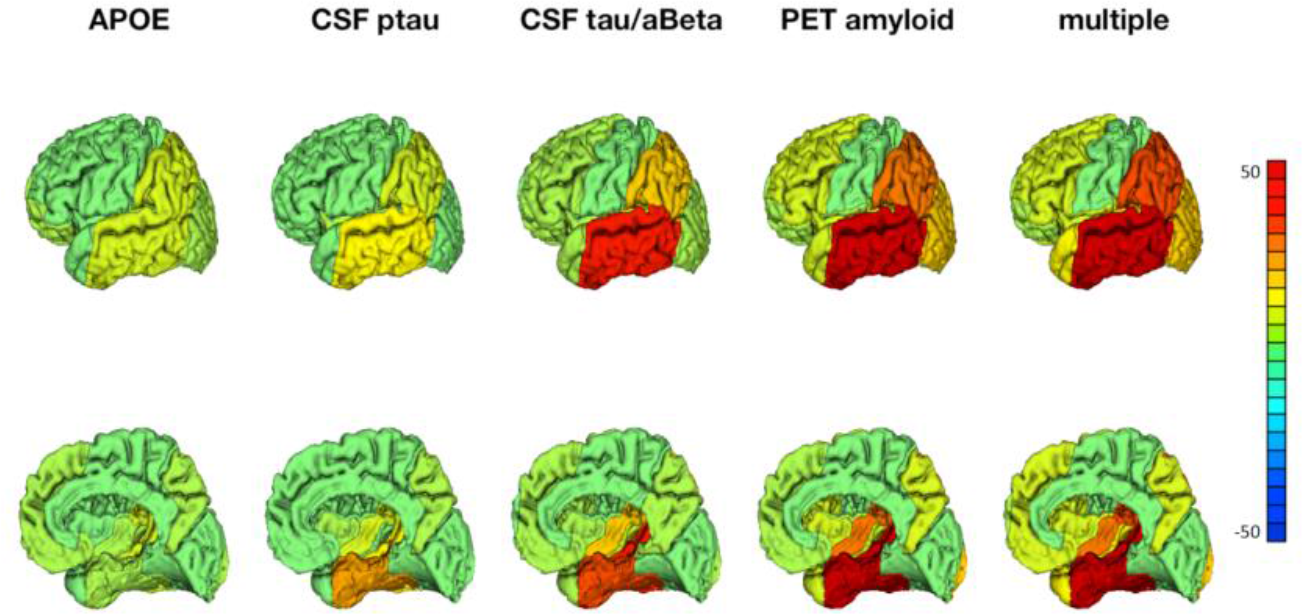
Log Bayes factors across cortex, comparing each biomarker-coupled MTL model to ‘*random*’.

**Figure 6.**
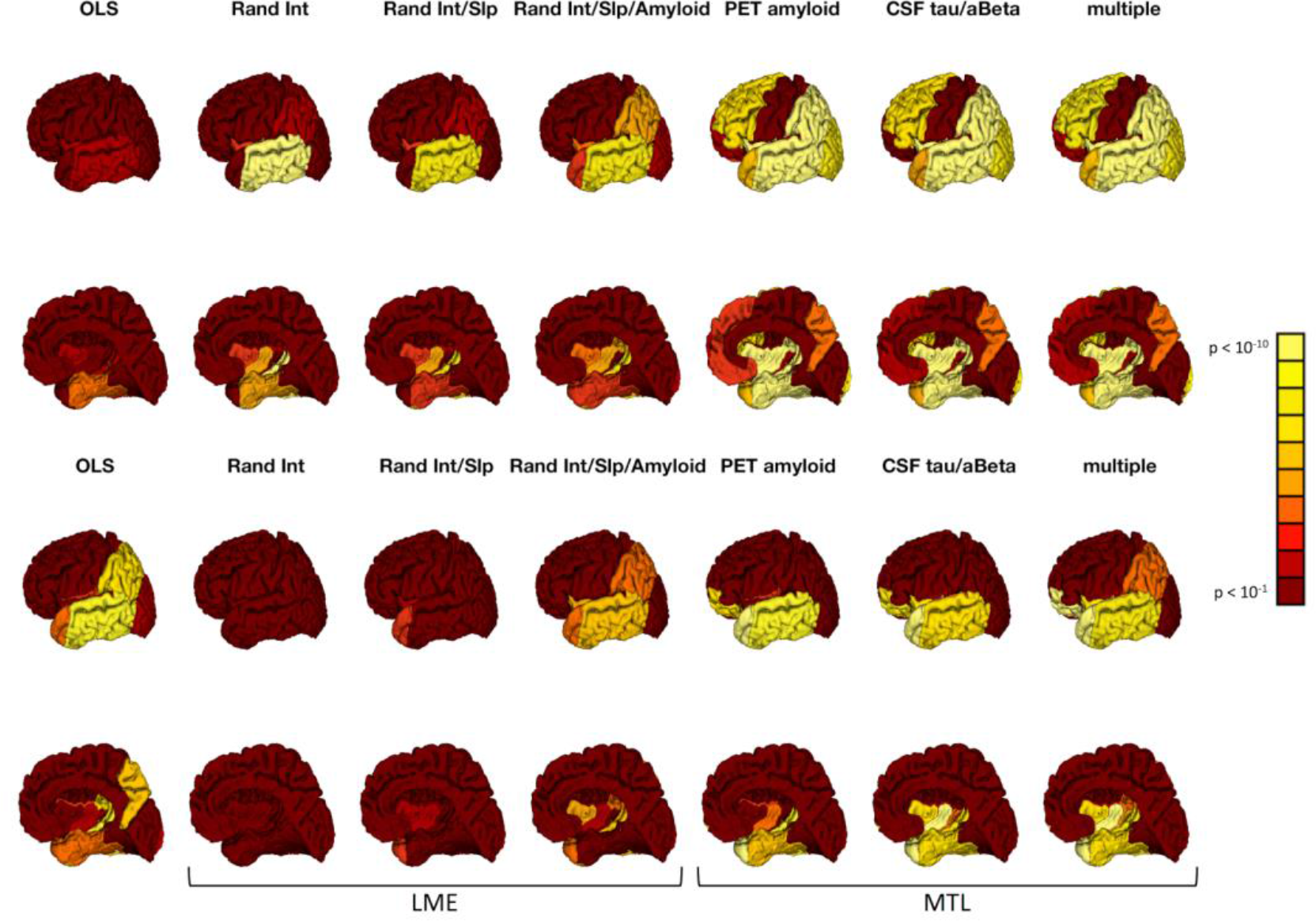
**Top:** Significance of (cross-sectional) diagnostic group differences in predicted volume at mean baseline age (73.5 years) for OLS, LME and selected MTL models **Bottom:** Same for (longitudinal) group differences in estimated slopes.

Longitudinally, the bottom row of Figure 6 shows both ‘*Rand Int*’ and ‘*Rand Int/Slp*’ have almost no significant slope differences in any region, while ‘*Rand Int/Slp/Amyloid*’ detects some significant differences within parts of the temporal lobe, insula and parietal regions, but does not detect the expected slope difference within the medial temporal lobe. The three MTL models, including ‘*PET amyloid*’, which represents the fairest comparison to the LME models, have significant differences across the temporal lobe (including the medial temporal lobe), insula, orbitofrontal region and, in the case of ‘*multiple*’, the lateral parietal region. Overall the MTL models infer a more plausible pattern of both cross-sectional and longitudinal disease effects than standard LME models.

It is reassuring that similar types of biological coupling (amyloid load measured via CSF and PET in ‘*CSF tau/aBeta*’ and ‘*PET amyloid*’ respectively) result in similar patterns of longitudinal differences. The longitudinal differences in the lateral parietal region detected by ‘*multiple*’ may be due to its incorporation of all biomarker coupling priors: ‘*APOE*’ and ‘*CSF ptau*’ also show some differences within that region (Figure S8, bottom row). In contrast to these biomarker-coupled models, ‘*random*’ does not detect any significant slope differences while ‘*OLS*’ detects few cross-sectional differences; neither model is implausible given other studies of AD-related atrophy (Frisoni et al., 2010; Risacher et al., 2010; Ziegler et al., 2015).

Interestingly, ‘*plain*’ only detects longitudinal differences within the medial temporal and temporal pole, suggesting that while this model can reliably detect strong true effects (see simulations), it may not be as sensitive as models with additional prior information. Further to this, Figure S9 depicts data for two regions: the medial temporal region, where most models agree that there are both cross-sectional and longitudinal differences, and the lateral temporal region, where ‘*plain*’ detects no longitudinal differences, though they are clearly evident in the figure. We further observe that ‘*APOE*’ detects a similar though weaker pattern of longitudinal differences compared to the other biomarker coupled models, suggesting that coupling based on similarity of genetic AD risk, conferred at birth, is less informative than coupling based on levels of amyloid accumulated decades later in older adults.

We also tested for cortical differences in subjects with differing numbers of APOE *ε*4 alleles (either 0, 1 or 2), analysing each diagnostic group separately. Figure S10 depicts group differences in number of alleles for ‘*APOE’*, ‘*PET amyloid*’ and ‘*multiple*’. ‘*OLS*’, ‘*plain*’ and ‘*random*’ showed no differences in neither baseline volumes nor slopes (data not shown) while ‘*CSF ptau*’ and ‘*CSF tau/aBeta*’ (not shown) had spatial patterns resembling that of ‘*PET amyloid*’, which shows allele-related differences in temporo-parieto-frontal, insular and anterior cingulate regions within the MCI group. There is a more widespread pattern of slope differences in ‘*APOE*’ and ‘*multiple*’ which is consistent with Ziegler et al., (2015), who found slope differences within temporo-parieto-frontal cortical grey matter in stable MCI subjects. However, these models also find slope differences within the insula and anterior cingulate in MCI subjects that were not reported in that study.

## Discussion

We have presented a multi-task learning based approach to modeling individuals’ longitudinal biomarker trajectories, setting the learning of each trajectory as a ‘task’ and using flexible covariance priors to couple tasks (i.e., subjects) during model training. Thanks to its parametric Bayesian formulation, our approach makes probabilistic predictions, infers distributions over parameters and allows for the comparison of competing models via model evidence. Using empirical Bayes (rather than time-consuming cross-validation), we showed how we can combine (i) fully decoupled models (i.e. individual-specific trajectory models; OLS-like); (ii) fully coupled models (i.e. a common trajectory across subjects; LME-like); and (iii) models coupled via one or more biomarker-based similarity matrices (i.e. kernels). In this way, our approach uses multi-kernel learning and capitalizes on different aspects of biology measured by different biomarkers, within a multi-task learning framework.

We performed simulations of trajectories having group wise variations in intercept and slope, showing that even the simplest version of our proposed model (‘*plain*’, mixing decoupled and fully coupled models) dramatically outperforms decoupled models (‘*OLS*’) in terms of predictive accuracy. We achieved further reductions in prediction error by adding kernel-based coupling using both random information and biomarkers (in various configurations, with and without oracle-like knowledge of simulation scenario). Interestingly, random-information-based kernels performed almost as well as the biomarker-based kernels (Figure 2), though the biomarker-based models (‘*Gaussian both*’ and the oracle-like models of each scenario) had better inference of true group differences (Figure 3). As such, biomarker-based models are the better choice for accurately making predictions and inferring parameters. We further conclude that ‘*Gaussian both*’, which allows biomarker-based coupling in all parameter types (e.g. intercepts and slopes) is a better choice than ‘*linear both*’ in terms of predictive performance and parameter inference. We emphasize the importance of parameter inference for both scientific (e.g. model interpretation) and translational purposes (e.g. trajectory parameter-based biomarkers such as ROI rates of change).

In this paper, we used empirical Bayes to estimate the coupling and noise hyperparameters, leading to a point estimate of the prior values of these variables. However, the full Bayesian approach may better account for both parameter and predictive uncertainty by placing priors on hyperparameters and estimating their posterior distributions. In our simulations, the two approaches had similar prediction errors and coverages. The models differed more in their parameter estimates: full Bayes had better parameter coverage and lower parameter error in some cases. On the other hand, empirical Bayes’ gradient descent based hyperparameter optimization runs significantly faster and scales better with the number of tasks than full Bayes’ MCMC sampling, thus providing a critical advantage of EB over full Bayes in real world applications. It is important to note though that in cases where the necessary posteriors can be derived, Gibbs sampling can significantly reduce this computational burden, making full Bayes more appealing (see e.g. Huertas et al., 2017).

We also tested the assumption of independent measurement noise, parameterized by β in Eq. (6), with an additional set of simulations. In general, prediction and parameter estimation errors were similar to or lower than LME models across varying noise correlations (Figure S2). However, some degradation of parameter coverage was evident, particularly at the highest noise correlation level, suggesting that a more general parameterization of the measurement error covariance (e.g. a block diagonal form allowing within-subject correlation) may be necessary in some situations. We did not explore the effect of non-Gaussian distributed measurement error on prediction and parameter inference here, although this may sometimes be an important consideration. In many cases, it may be possible to address this violation using published approaches (e.g. variable transformation).

We applied the model to longitudinal data from the ADNI study, modeling trajectories of cortical ROIs across CN, MCI and AD subjects using kernels formed from amyloid PET, CSF and genetic (APOE) information. We showed degeneration throughout the cortex, with lateral regions degenerating faster than medial regions (Figure 4). We showed significantly decreased prediction errors due to coupling (‘*plain*’ versus *‘OLS*’) and further decreases when adding kernel-based coupling, with a small difference between the random-information versus biomarker coupled models in some ROIs (Figure 4 and Figure S6).

Our model offers improved interpretability and more concrete biological explanations of trajectory differences across diagnostic groups compared to the baseline models. Here, we were mainly interested in understanding how cortical degeneration varies across diagnostic groups, which required that we carefully interpret the patterns of group differences (Figures 6 and S8). Single biomarker models based on either ‘*PET amyloid*’ or ‘*CSF tau/aBeta*’ had cross-sectional and longitudinal group differences that were consistent with the literature and had the most evidence in their favor (Figure 5). Importantly, this analysis showed the benefit in coupling cortical trajectories based on baseline measures of amyloid deposition measured via PET or amyloid-to-tau ratio via CSF, which is consistent with the prevailing disease progression model of AD in which amyloid deposition precedes change in brain structure (Jack et al., 2010). Coupling based on genetic risk for AD as realized by the APOE genotype was inferior to using baseline amyloid-based biomarkers. This agrees with the literature in that APOE genotype is the genetic risk and amyloid biomarker levels represent the realization of that risk. Furthermore, we showed that combining multiple kernels is effective in the sense that ‘*multiple*’, the multi-modal model, was as good as the best individual model in terms of model evidence and parameter inference. Thus, our approach removes the requirement to pre-select any specific biomarker.

All coupled models had significant diagnostic group differences across the cortex at mean baseline age, agreeing with the pattern of later-stage neurofibrillary changes due to AD that have been shown to be detectable via MRI (Braak and Braak, 1991; Whitwell et al., 2008), along with many of the AD discrimination studies that have used cross-sectional structural MRI based features (Arbabshirani et al., 2016). In particular, the pattern of cross-sectional differences we find aligns with Karas et al., (2003), which reported AD-related differences within the temporal lobe and insula, with sparing of the sensorimotor cortex. We also found no significant differences within the motor and sensory ROIs, supporting the idea that sensorimotor function is relatively spared in AD, unless the disease is very advanced (Suvà et al., 1999; Ferreri et al., 2016). Among the models with high model evidence in their favor, namely ‘*CSF tau/aBeta*’, ‘*PET amyloid*’ and ‘*multiple*’, there were significant longitudinal (i.e. slope) differences within the temporal lobe, orbitofrontal region, insula and lateral parietal region. These findings are similar to the patterns of group differences in one year atrophy depicted in Risacher et al., (2010), although the authors did not focus on their apparent findings within the insula. Insel et al., (2015) identified changes within the insula and temporal regions occurring prior to the clinical threshold for amyloid-β positivity, and interestingly, we detect longitudinal differences in these regions with models that couple based on similarity of protein measures.

In addition to clinical diagnosis, we also investigated the effect of APOE ε4, the major genetic risk factor for late-onset AD, analysing differences in subjects grouped by number of ε4 alleles (Figure S10). We found no cross-sectional volume differences at mean baseline age within each group and few significant longitudinal differences within the CN and AD groups. The CN finding is consistent with the literature: Filippini et al., (2009) found no volumetric differences within the brain between young, healthy ε4 carriers and matched non-carriers using cross-sectional information while Raz et al., (2010) found no differences due to ε4 within healthy middle-aged and older adults using longitudinal data. Our findings indicate similar homogeneity within AD subjects. Within the MCI group we found a temporo-parietal-frontal pattern of slope differences that aligned with previous literature (Ziegler et al., 2015) along with additional slope differences with the insula and anterior cingulate. We note that the findings within this group may be due to both the larger sample size and greater heterogeneity of the MCI group compared to the CN and AD groups.

We also compared our novel MTL approach to a widely available LME implementation, showing that MTL makes very similar prediction errors on the held-out ADNI follow-ups (Figure S7). However, MTL detected more widespread cross-sectional group differences than the three LME models we considered and, importantly, more significant longitudinal differences within the temporal lobe (Figure 6). As such the MTL based parameters appear to be more plausible than the LME based parameters. Additionally, our method automatically finds the covariance structure that best explains the training data (within the limit of the chosen parameterization), removing modeling decisions such as whether a variable is or is not a random effect.

The approach we have presented has several limitations however. Firstly, computing the log marginal likelihood at each optimization step involves the inversion of the prior covariance matrix (see Appendix), which scales cubically with the number of subjects in the worst case. This precludes coupling beyond hundreds of subjects and restricts us to univariate modeling. A multivariate approach would exacerbate the problem, scaling cubically with the product of subjects by variables. Reduced rank approaches or inducing point methods may speed up computation, as would a diagonal approximation of the matrix inversions, sacrificing accuracy for speed. Alternatively, one could use GPUs; Tensorflow has highly optimised linear algebra routines for matrix operations that can deliver an order of magnitude speed improvements (Abadi et al., 2016). Secondly, beyond computational considerations, our model may not capture long term, nonlinear trends that are only evident across subjects (see e.g. Donohue et al., (2014)). To properly model these may require adding a fixed effects component to accommodate higher-order polynomial functions describing group-level trajectories. Alternatively, one may switch to modeling trajectories of study time, which may introduce significant intercept differences.

There are multiple directions for future work. As mentioned in the introduction, the method we present is not a disease progression model and as such it does not estimate a disease stage for each subject. It does, however, provide plausible estimates of trajectory parameters which may serve as valuable inputs to a staging model. Future work will investigate the staging of subjects based on these parameters within an EBM (Young et al., 2014), providing insight into the role of brain structural changes during the progression from normal cognition to Alzheimer’s disease (Jack et al., 2010). We can also extend our understanding of the relationship between genetics and cortical atrophy beyond APOE to all single nucleotide polymorphisms (SNPs) using multivariate methods such as partial least squares (PLS; Lorenzi et al., 2018) or canonical correlation analysis (Szefer et al., 2017). Finally, we can generalize the approach to simultaneously model multiple variables across subjects (i.e. multi-output learning), where interesting modeling possibilities (coupling parameters across variables within and between subjects) and computational challenges abound. Note, however, that some studies have shown only a modest benefit of this type of coupling relative to the computational effort involved (Zhang and Shen, 2012; Marquand et al., 2014).

## Acknowledgements

Authors are grateful to Dr. Marco Lorenzi and Prof. Jonathan Schott for their comments and suggestions. This project has received funding from the European Union’s Horizon 2020 research and innovation programme under grant agreement No. 666992. EPSRC grants M020533, M006093, J020990 additionally support DCA’s work on this topic. Andre Altmann holds an MRC eMedLab Medical Bioinformatics Career Development Fellowship. This work was supported by the Medical Research Council [grant number MR/L016311/1]. Marzia Antonella Scelsi acknowledges financial support by the EPSRC-funded UCL Centre for Doctoral Training in Medical Imaging (EP/L016478/1). Andre Marquand gratefully acknowledges support from the Netherlands Organisation for Scientific Research (NOW) via a Verniewingsimpuls VIDI fellowship (91716415). Sebastien Ourselin receives funding from the EPSRC (EP/H046410/1, EP/J020990/1, EP/K005278), the MRC (MR/J01107X/1), the EU-FP7 project VPH-DARE@IT (FP7-ICT-2011-9-601055), the NIHR Biomedical Research Unit (Dementia) at UCL and the National Institute for Health Research University College London Hospitals Biomedical Research Centre (NIHR BRC UCLH/UCL High Impact Initiative-BW.mn.BRC10269).

Data collection and sharing for this project was funded by the Alzheimer’s Disease Neuroimaging Initiative (ADNI) (National Institutes of Health Grant U01 AG024904) and DOD ADNI (Department of Defense award number W81XWH-12-2-0012). ADNI is funded by the National Institute on Aging, the National Institute of Biomedical Imaging and Bioengineering, and through generous contributions from the following: AbbVie, Alzheimer’s Association; Alzheimer’s Drug Discovery Foundation; Araclon Biotech; BioClinica, Inc.; Biogen; Bristol-Myers Squibb Company; CereSpir, Inc.; Cogstate; Eisai Inc.; Elan Pharmaceuticals, Inc.; Eli Lilly and Company; EuroImmun; F. Hoffmann-La Roche Ltd and its affiliated company Genentech, Inc.; Fujirebio; GE Healthcare; IXICO Ltd.; Janssen Alzheimer Immunotherapy Research & Development, LLC.; Johnson & Johnson Pharmaceutical Research & Development LLC.; Lumosity; Lundbeck; Merck & Co., Inc.; Meso Scale Diagnostics, LLC.; NeuroRx Research; Neurotrack Technologies; Novartis Pharmaceuticals Corporation; Pfizer Inc.; Piramal Imaging; Servier; Takeda Pharmaceutical Company; and Transition Therapeutics. The Canadian Institutes of Health Research is providing funds to support ADNI clinical sites in Canada. Private sector contributions are facilitated by the Foundation for the National Institutes of Health (www.fnih.org). The grantee organization is the Northern California Institute for Research and Education, and the study is coordinated by the Alzheimer’s Therapeutic Research Institute at the University of Southern California. ADNI data are disseminated by the Laboratory for Neuro Imaging at the University of Southern California.

## Appendix

We wish to find the hyperparameters that maximize the model’s marginal likelihood. For computational and analytic reasons, it is easier to minimize the negative (natural) log marginal likelihood, setting this as the optimizer’s objective. We need the partial derivatives with respect to each hyperparameter, constraining each to be strictly positive. For the noise term β this is a natural constraint; for the coupling weights **α** we follow the rule that a positive sum of valid kernels is also a valid kernel. To impose these constraints within an unconstrained optimizer we transform the variables, optimizing log(β) and log(**α**), which will be strictly positive when exponentiated. To derive the necessary partial derivatives with respect to the transformed variables we make use of the chain rule:

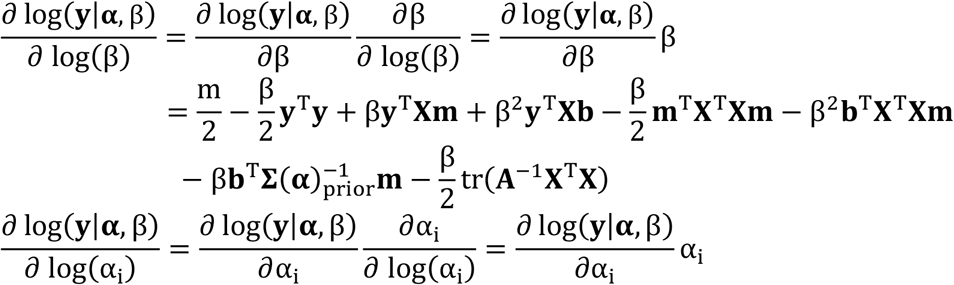

where α_i_ is an element within vector **α**, **b** = (**I**_nd_ − β**A**^−1^**X**^T^**X**)**A**^−1^**X**^T^**y** and

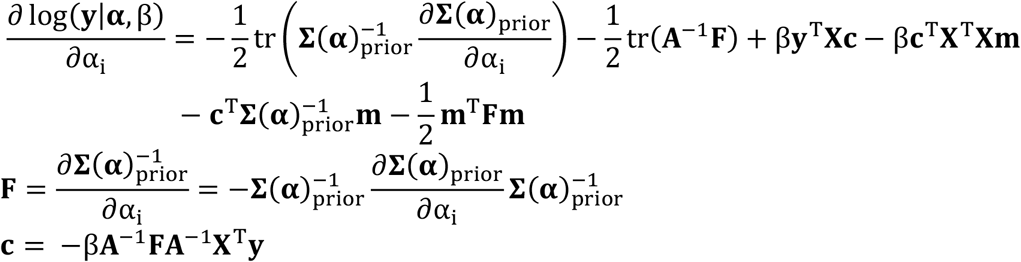

and the ∂**Σ**(**α**)_prior_/∂α_i_ depends on the α_i_ in question. We can easily change the parameterization of **Σ**(**α**)_prior_prior without breaking these equations, provided it remains invertible and differentiable with respect to its parameters.

For the form used in Eq.’s (3) - (5), we need ∂**Σ**(**α**)_prior_/∂α_1_ = **I**_nd_, ∂**Σ**(**α**)_prior_/∂α_i1_ = **I**_n_**⊗M**_ii_, ∂**Σ**(**α**)_prior_/∂α_i2_ = **1**_n_**⊗M**_ii_ and ∂**Σ**(**α**)_prior_/∂α_i(j+2)_ = **K**_j_**⊗M**_ii_ for the matrix weighting hyperparameters, where i = 1, …, d and j = 1, …, k. Some kernels may also have internal hyperparameters (also included in **α**); one such example is the σ_SE_ parameter of the SE kernel. In this case, we need ∂**Σ**(**α**)_prior_/∂σ_SE_ = −α_i(j+2)_(**D⊙K**_j_)**⊗M**_ii_ where **K**_j_ = exp(−σ_SE_**D**) and ⊙ is the element-wise product.

## Footnotes

1 In practice, we include an additional diagonal term *ε***I**_nd_, with *ε* set to 10^−6^ throughout, that aids numerical stability when inverting **Σ**_prior_.

2 There may be more hyperparameters if the kernels themselves contain tuneable parameters.

3 We do not vary the biomarker noise here as we found that, in general, it did not have a strong effect on the models, particularly as compared to the measurement noise.

4 https://surfer.nmr.mgh.harvard.edu/fswiki/LinearMixedEffectsModels

