## Supplementary material for "Modeling longitudinal imaging biomarkers with parametric Bayesian multi-task learning": Table S1

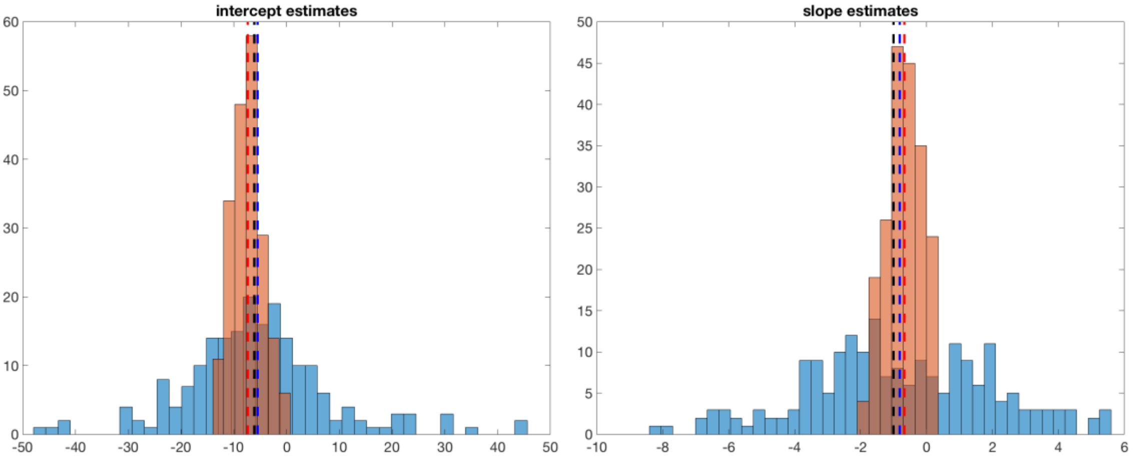


**Figure S1 Left:** histogram of intercept estimates from a representative MTL model (‘*plain*’) in red versus ordinary least squares’ estimates in blue, with corresponding red and blue dashed lines indicating each model’s mean estimates and black line indicating the true mean across intercepts. **Right:** same for slope estimates.

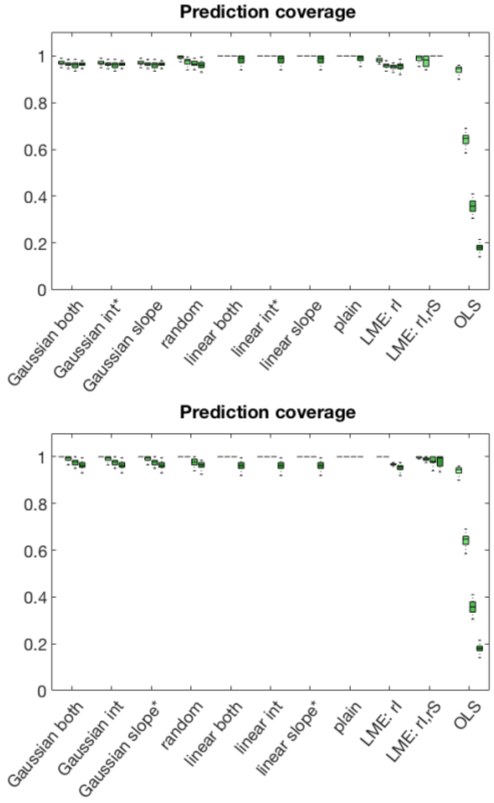


**Figure S2** Boxplots of models’ prediction coverage probabilities for intercept variation (top figure) and slope variation (bottom figure) simulations.

**Table S1** Convergence diagnostics for the hyperparameters of the '*MCMC plain*' model for one representative run of the intercept varying simulation scenario. MCSE is the Markov Chain standard error, defined as the standard deviation ($\sigma$) divided by $\sqrt{N_{eff}}$, where $N_{eff}$ is effective sample size.

| Name | $\mu$ | $\sigma$ | $N_{eff}$ | $\hat{R}$ | MCSE | Confidence Interval |
|---|---|---|---|---|---|---|
| $\alpha_1$ | 0.032 | 0.032 | 479 | 1 | 0.0014 | (0.029, 0.035) |
| $\alpha_{11}$ | 4.9 | 1.4 | 4000 | 1 | 0.022 | (4.85, 4.94) |
| $\alpha_{12}$ | 75 | 56 | 4000 | 1 | 0.89 | (73.3, 76.7) |
| $\alpha_{21}$ | 0.033 | 0.033 | 459 | 1 | 0.0015 | (0.030, 0.036) |
| $\alpha_{22}$ | 55 | 52 | 4000 | 1 | 0.83 | (53.4, 56.6) |
| $\beta^{-1}$ | 1.2 | 0.24 | 1539 | 1 | 0.0062 | (1.19, 1.21) |

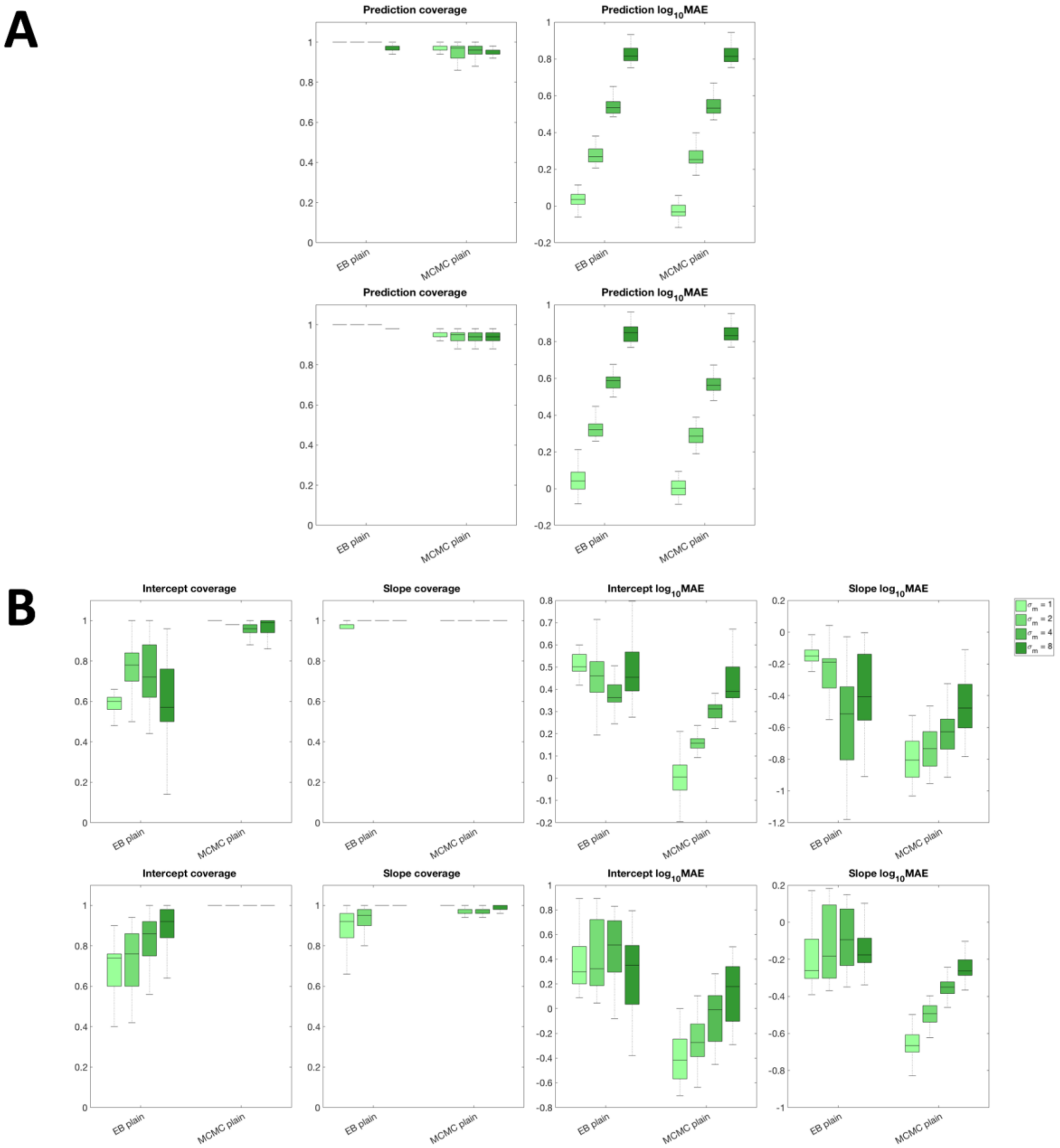


**Figure S3** Simulations comparing proposed empirical Bayesian realization of ‘*plain*’ (‘*EB plain*’) to its full Bayesian realization via MCMC sampling (‘*MCMC plain*’). **A:** boxplots of prediction errors ($\log_{10}$MAE) and prediction coverage probabilities for both models. Top row is intercept varying scenario, bottom row is slope varying scenario. **B:** corresponding boxplots of parameter coverage and parameter prediction error for both scenarios.

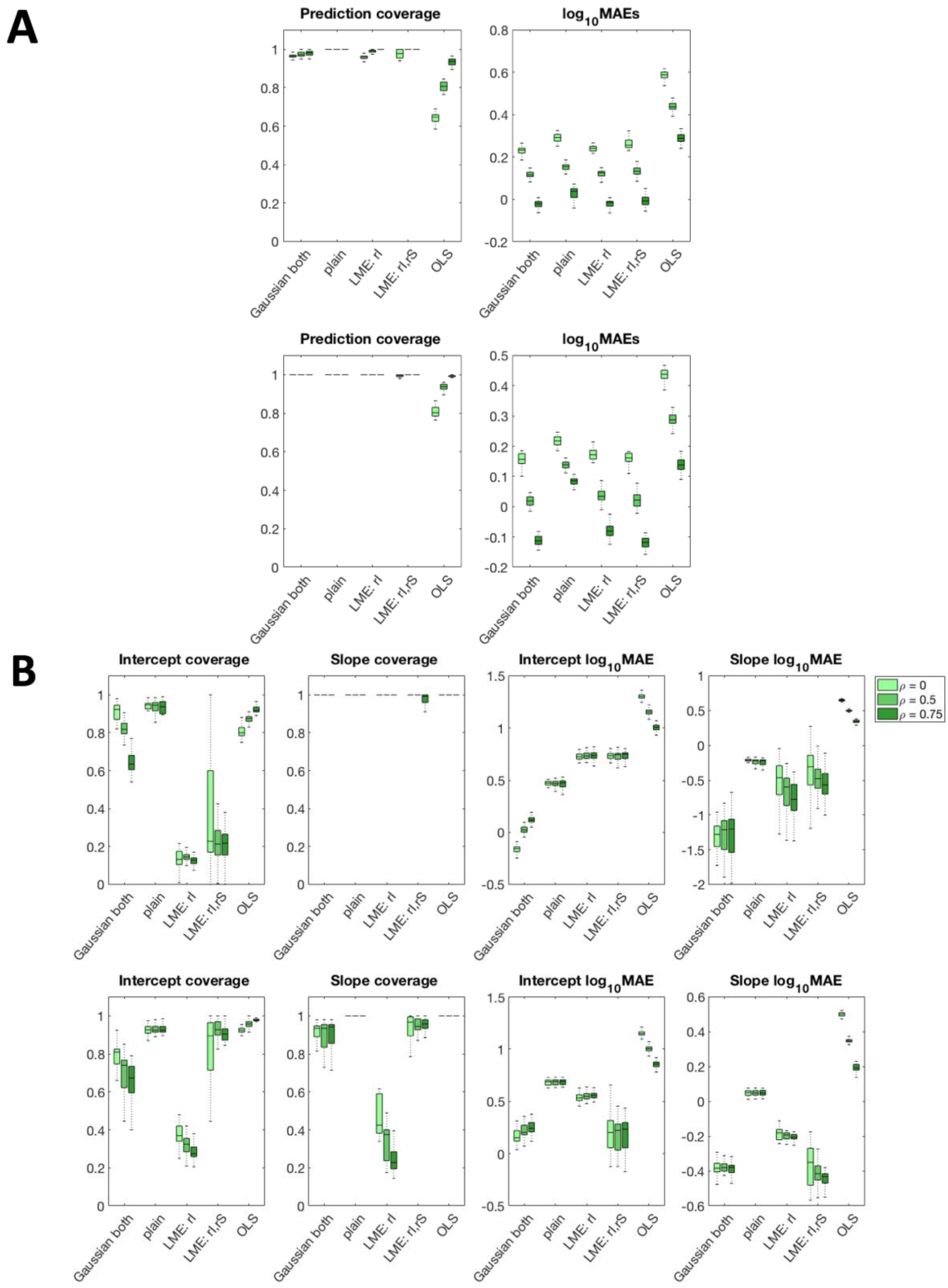


**Figure S4** **A:** boxplots of prediction errors ($\log_{10}$MAE) and prediction coverage probabilities for simulations varying measurement noise correlation (parameter $\rho$) for four representative models. Top row is intercept varying scenario, bottom row is slope varying scenario. **B:** corresponding boxplots of parameter coverage and parameter prediction error for both scenarios.

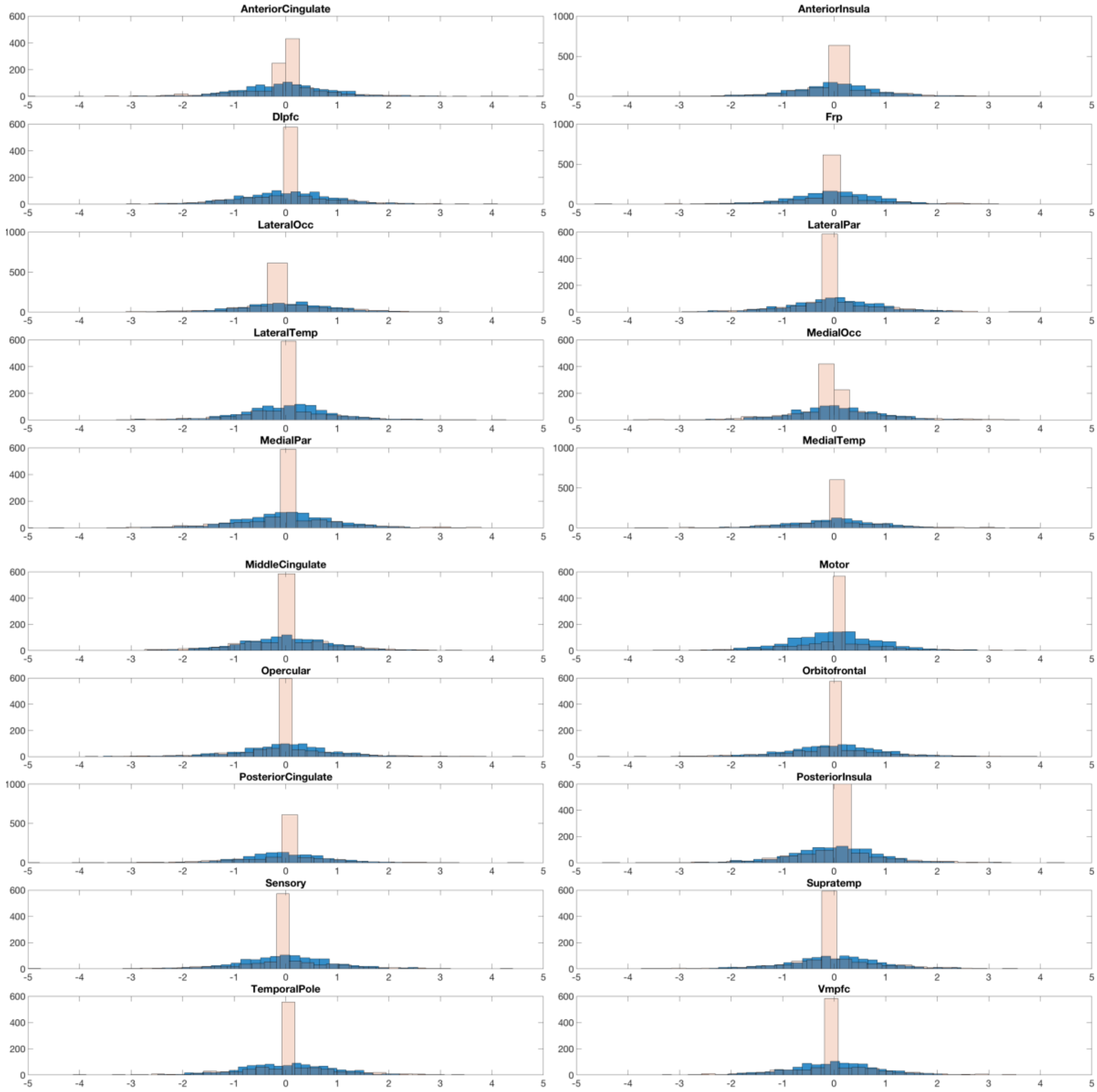


**Figure S5** Histograms of residuals for the '*CSF tau/aBeta*' (blue) and '*OLS*' (rose) models for each region of interest in the ADNI application.

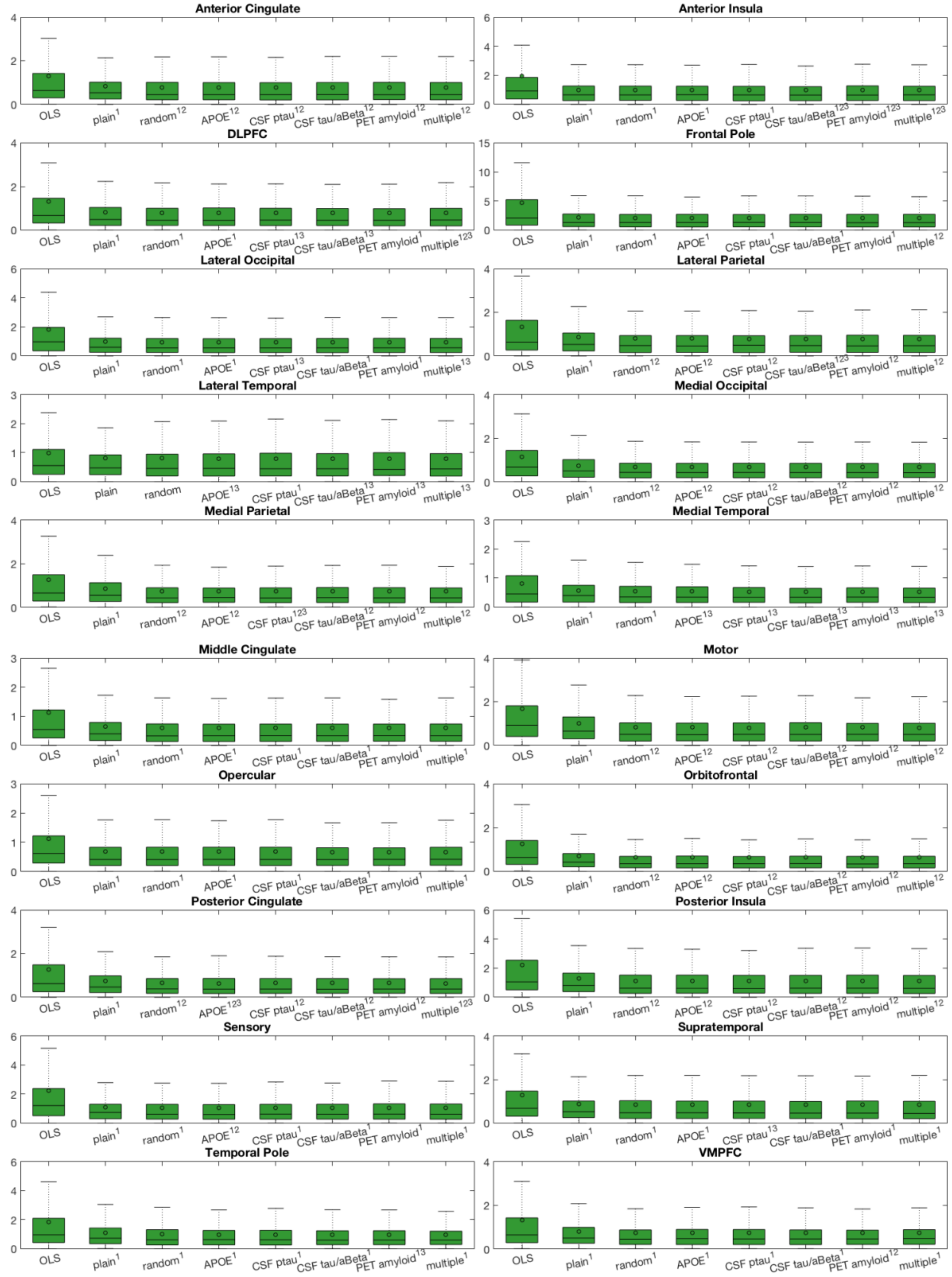


**Figure S6** Boxplots (plus mean value as circle) of absolute errors between actual and predicted annualized rate of change from baseline to final (out-of-sample) follow-up (y-axis), across all models (x-axis) and cortical ROIs (panels). Superscripts: significantly lower MAE ($p < 0.05$) of given model compared to 1: '*OLS*'; 2: '*plain'*; 3: '*random'*. Abbreviations: DLPFC: dorsolateral prefrontal cortex; VMPFC: ventromedial prefrontal cortex.

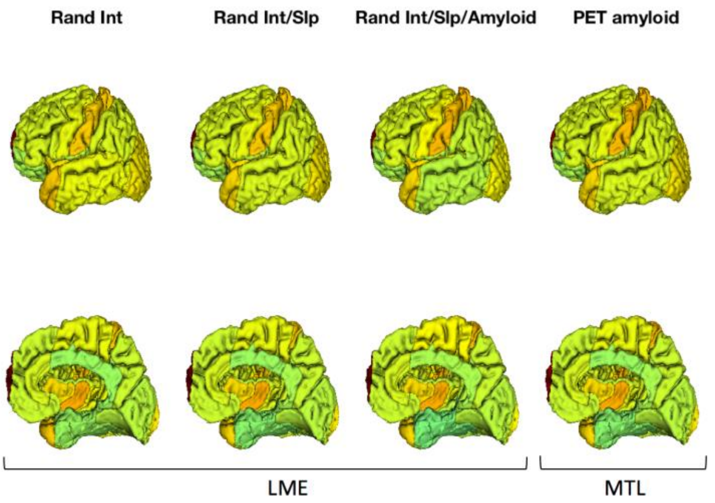


**Figure S7** Prediction errors (MAEs of annualized rates of change) for three LME models and most comparable MTL model (‘*PET amyloid*’).

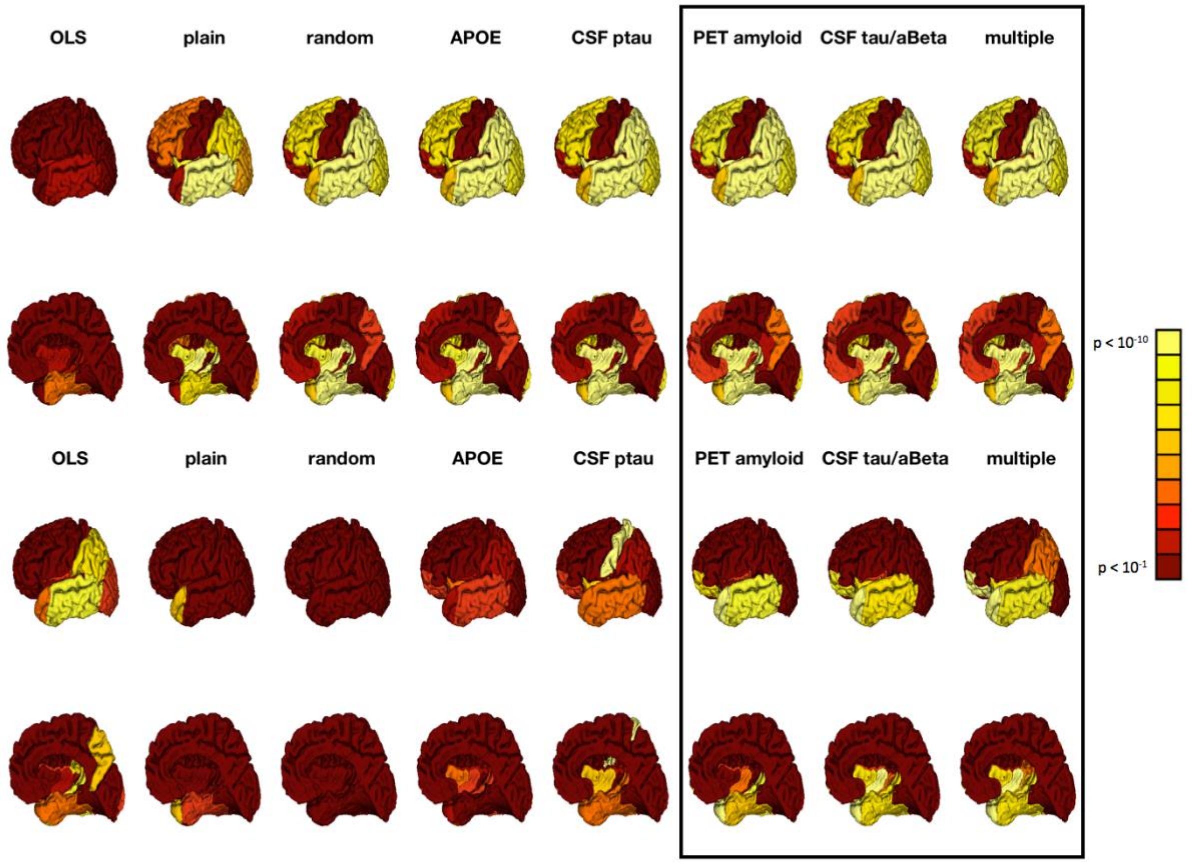


**Figure S8 Top:** Significance of (cross-sectional) diagnostic group differences in predicted volume at mean baseline age (73.5 years) across cortex for all MTL models **Bottom:** Same for (longitudinal) group differences in estimated slopes across all MTL models. ‘*CSF tau/aBeta*’, ‘*PET amyloid*’, ‘*multiple*’ have largest model evidence relative to ‘*random*’ (see Figure 5).

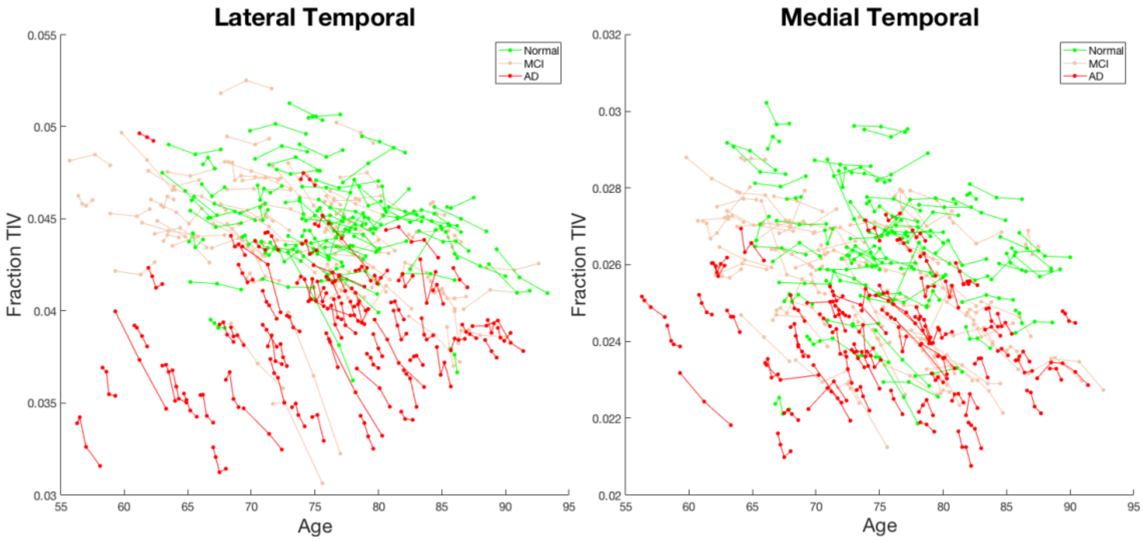


**Figure S9** All available data-points for seventy randomly selected subjects per group in two regions (210 subjects total), showing both (cross-sectional) group differences at mean age (73.5 years) and (longitudinal) differences in rates of change.

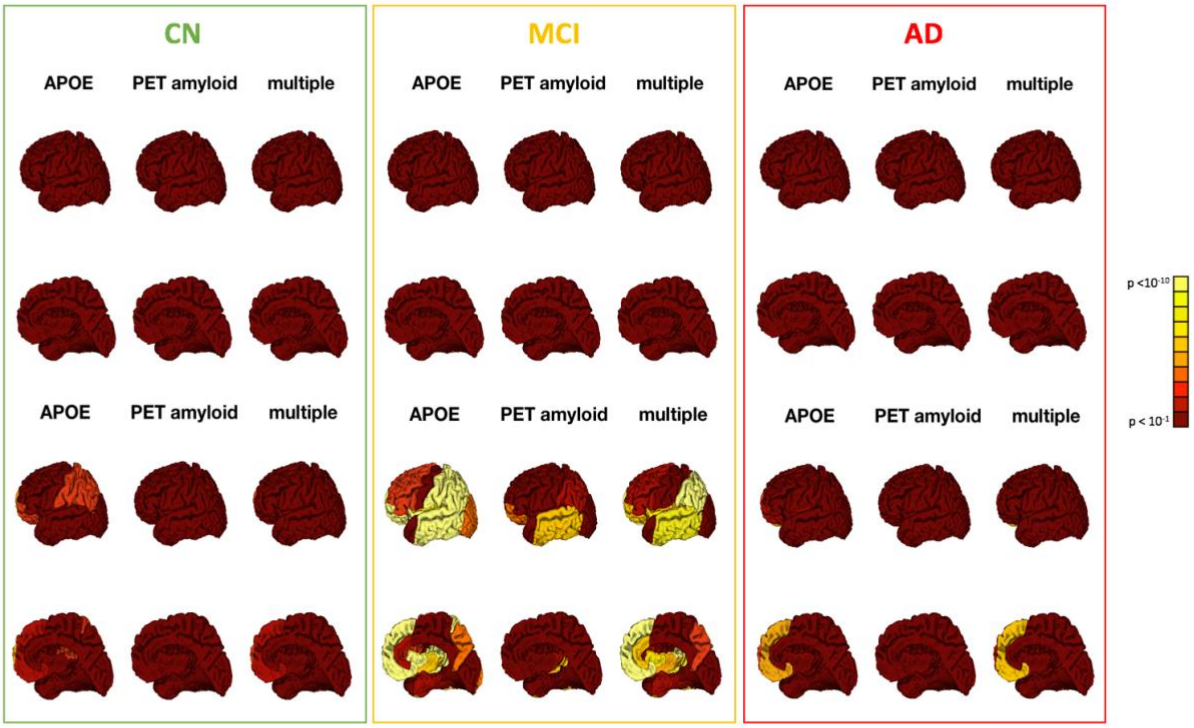


**Figure S10 Top:** Effect of the number of APOE ε4 alleles on cortical volume at mean baseline age (73.5 years) within each diagnostic group for three representative models, Bonferroni corrected for all comparisons **Bottom:** Same for effect of number of alleles on estimated slopes.
